# Dual DNA-binding capability of Cdc13 coordinates with Ku to safeguard telomere integrity

**DOI:** 10.1101/2025.09.02.671137

**Authors:** Zhitong Feng, Jiangchuan Shen, Yuxi Li, Giovanni Gonzalez-Gutierrez, Quan Wang, Hengyao Niu

## Abstract

In eukaryotes, telomeres, nucleoprotein complexes assembled on telomeric 3’-overhang DNA, protect linear chromosome ends from degradation and inappropriate DNA repair. In budding yeast, *Saccharomyces cerevisiae,* telomere assembly involves the Cdc13-Stn1-Ten1 (CST) complex, the Ku70-Ku80 (Ku) complex, and the Rap1-Rif1-Rif2 complex. Among these, the CST complex, centered on the binding of the G-rich single-stranded region by the Cdc13 subunit, is essential for telomere maintenance and protection. Here, we show that Cdc13 also interacts with the duplex DNA adjoining its bound single strand. This junction binding enables Cdc13 to reposition the Ku complex, potentially suppressing Ku-mediated end joining, while allowing the Cdc13 and Ku complexes to synergize in protecting telomeric DNA ends, a function that can be disrupted by mutation of a conserved residue (K504E). *cdc13-K504E* cells remain viable but are hypersensitive to Exo1 overexpression, particularly when combined with *ku80Δ*. Surprisingly, *cdc13-K504E* and other telomere protection mutants are prone to adaptive metabolic reprogramming during stationary-phase growth, leading to increased fitness. This reprogramming is inheritable and centers on elevated expression of thiamine biosynthesis, salvage, and intake pathways. Our findings reveal a stress-induced adaptive response associated with telomere erosion, indicating that telomere erosion may serve as a timer for its activation.

## Introduction

Telomeres are specialized nucleoprotein structures that cap the ends of linear chromosomes and are essential for preserving genome integrity. They prevent chromosome ends from being recognized as double-strand breaks (DSBs), thereby avoiding aberrant repair events such as end-to-end fusion, translocation, and genome instability^1–3^. In eukaryotes, telomeres typically consist of repetitive G-rich DNA ending in a 3′ single-stranded overhang, which in humans spans several kilobases of TTAGGG repeats, and in *Saccharomyces cerevisiae* consists of ∼300 bp dsDNA and a 12–15 nt TG₁–₃ 3′ overhang^1,3,4^.

Despite their protective role, telomeres pose a unique paradox: they must simultaneously suppress DNA damage responses while still permitting telomeric DNA ends processing, which is essential for telomere maturation^2,3^. This dilemma is most evident during S phase, when telomeric ends undergo maturation involving limited nucleolytic resection and telomerase catalyzed 3’ end extension^4,5^. In budding yeast, the same nucleolytic resection machinery, essential for telomere maturation, which generates the 3’ overhang required for telomerase action, is also central to homologous recombination (HR) repair of double-strand breaks (DSBs)^6–9^. This DNA end resection is primarily executed by three conserved pathways: the MRX complex (Mre11-Rad50- Xrs2) with Sae2, the exonuclease Exo1, and the helicase-nuclease ensemble Sgs1–Dna2^6,7,10–12^. Notably, Exo1 and Sgs1–Dna2 require a free DNA end to initiate resection, whereas MRX–Sae2 can process blocked or internal ends^7,12^. While limited resection is indispensable for telomere maintenance, uncontrolled or aberrant resection poses a significant threat to telomere integrity^4,10,13^. This inherent conflict underscores why restraining resection is critical for telomere protection. To resolve this paradox, cells have evolved “telomere capping” complexes (e.g., shelterin in mammals)^2^ that preserve the telomere’s integrity while regulating resection/repair enzymes’ access to chromosomal ends^2,3,9^.

In budding yeast, telomere capping involves three major modules: the Ku70/80 heterodimer, the Rap1–Rif1/2 complex, and the CST complex (Cdc13–Stn1–Ten1)^4,14^. Rap1 and its associated factors, Rif1/2, occupy the telomeric dsDNA region, regulate telomere length and subtelomeric silencing^15,16^. The Ku heterodimer also encircles telomeric dsDNA and protects against 5′ exonucleolytic degradation^17–21^, although Ku is more widely found at the ss/dsDNA junction of double-strand breaks (DSBs) to promote non-homologous end joining (NHEJ)^22,23^. Central to end protection is the CST complex, whose core component Cdc13 binds the 3′ overhang with high affinity. Structurally, Cdc13 contains four oligonucleotide/oligosaccharide-binding (OB) folds (OB1, 2, 3 and 4) and a telomerase recruitment domain (RD)^24,25^ adjoining OB1 and OB2. The N- terminal OB1 mediates homodimerization with auxiliary ssDNA-binding capacity, while OB3 harbors the core telomeric G-strand recognition activity (**Fig. S1A**) ^26,27^. This ssDNA engagement is essential for yeast viability, shielding telomeric termini from degradation while facilitating telomerase recruitment and C-strand fill-in for telomere maturation ^28,29^.

Notably, the CST complex demonstrates significant evolutionary conservation and functional divergence. In humans, CST plays essential roles in telomere replication and maturation, thereby contributing to telomere stability. It also participates in genome-wide replication regulation.^30–32^. Defects in CST components are associated with several human disorders, including Coats plus syndrome and dyskeratosis congenita, characterized by telomere dysfunction, genome instability, and cancer predisposition^33–37^. Aberrant CST expression is increasingly linked to tumorigenesis, underscoring the importance of understanding its molecular mechanisms.

Despite these advances, a fundamental question persists in the yeast model: How does Ku, a core NHEJ factor dedicated to end joining^17^, persistently localize to telomeres without catalyzing catastrophic end-to-end fusions? This perplexing behavior implies specialized mechanisms actively suppress Ku’s repair activity at chromosome ends. Yet, the mechanisms of silencing this potentially deleterious repair activity at chromosome ends remain poorly understood.

Here, we uncover a novel DNA-recognition interface of Cdc13 that enables it to not only bind telomeric ssDNA but also specifically recognize the ss/dsDNA junctions^38^. We show that this junction recognition is essential for Cdc13 to suppress both Exo1- and Sgs1–Dna2-mediated resection and to facilitate Ku repositioning from ss/dsDNA junctions. Unlike *cdc13Δ*, mutation that impairs junction recognition doesn’t cause lethality, nor telomere shortening. However, it sensitizes the cells to resection stress in a manner that is synergistic with Ku mutations. Interestingly, the junction binding mutant of Cdc13 led to the arising of colonies with greater fitness during the stationary phase growth and, to our surprise, the fitness gain can be carried to vegetative growth. FACS analysis suggested these colonies have faster cycling with an increase in the G2 population. Overall, our findings suggest Cdc13 may modulate Ku activity via its junction recognition to likely suppress Ku mediated end-joining but synergize with Ku in counteracting end resection stress at telomeres at the same time. The loss of this junction recognition, while not essential for telomere maintenance during vegetative growth, promotes adaptive changes of cells in the stationary phase. These insights bridge fundamental telomere biology with mechanisms underlying age-related disorders and cancer.

## Results

### Cdc13 interacts with ss/dsDNA junctions with enhanced affinity for telomeric 3′ overhangs

Cdc13 protects telomeric DNA largely through its DNA binding. To investigate its role in telomere protection, we first examined the substrate specificity of Cdc13 in telomeric DNA binding. To do so, recombinant full-length Cdc13 protein with an N-terminal SUMO tag was expressed in *E. coli* and purified to near homogeneity (**Fig. S1B**). After SUMO tag removal, size-exclusion chromatography (SEC) analysis indicated the apparent molecular weight as around 280 kDa consistent with its dimer formation as reported (**Fig. S1C**)^27^.

To characterize the DNA-binding activity and specificity of the purified Cdc13 protein, we designed a panel of DNA substrates with identical duplex regions but varying 3′ overhangs to mimic different telomeric contexts: Tel14 (14-nt TG₁-₃ overhang) reflecting the physiological range of mature yeast telomeres^4^; Tel22 representing an extended overhang optimal for engaging two Cdc13 molecules, based on prior studies showing that each Cdc13 binds a minimum of ∼11 nt^39,40^; and Tel7 (truncated 7-nt TG₁-₃ overhang) serving as a minimal-length control. In addition, a non-telomeric poly-dT overhang of 22 nt (T22) was included as a sequence specificity control. Electrophoretic mobility shift assays (EMSAs) under physiological salt conditions (150 mM KCl) revealed that Cdc13 binds Tel14 and Tel22 with high affinity, exhibiting distinct, cooperative banding patterns and apparent Kd values of ∼5 nM and <2.5 nM, respectively, indicative of tight interactions (**Fig. 1A (i) and (ii)**). In contrast, the non-telomeric poly-dT substrate T22 showed much weaker binding (Kd ∼80–100 nM) (**Fig. 1B(i)**), underscoring the sequence specificity of Cdc13 for telomeric G-rich ssDNA. Surprisingly, we also observed weak binding with Tel7 (apparent Kd ∼20 nM) (**Fig. 1A (iii)**), in stark contrast to its negligible binding to 7-nt ssDNA (ssTel7, **Fig. 1B (ii**)). This disparity aligns with prior reports that Cdc13 requires ≥11 nt for stable ssDNA binding^39,40^ and indicates that the duplex junction may compensate for suboptimal overhang length by providing additional contacts.

**Figure 1.**
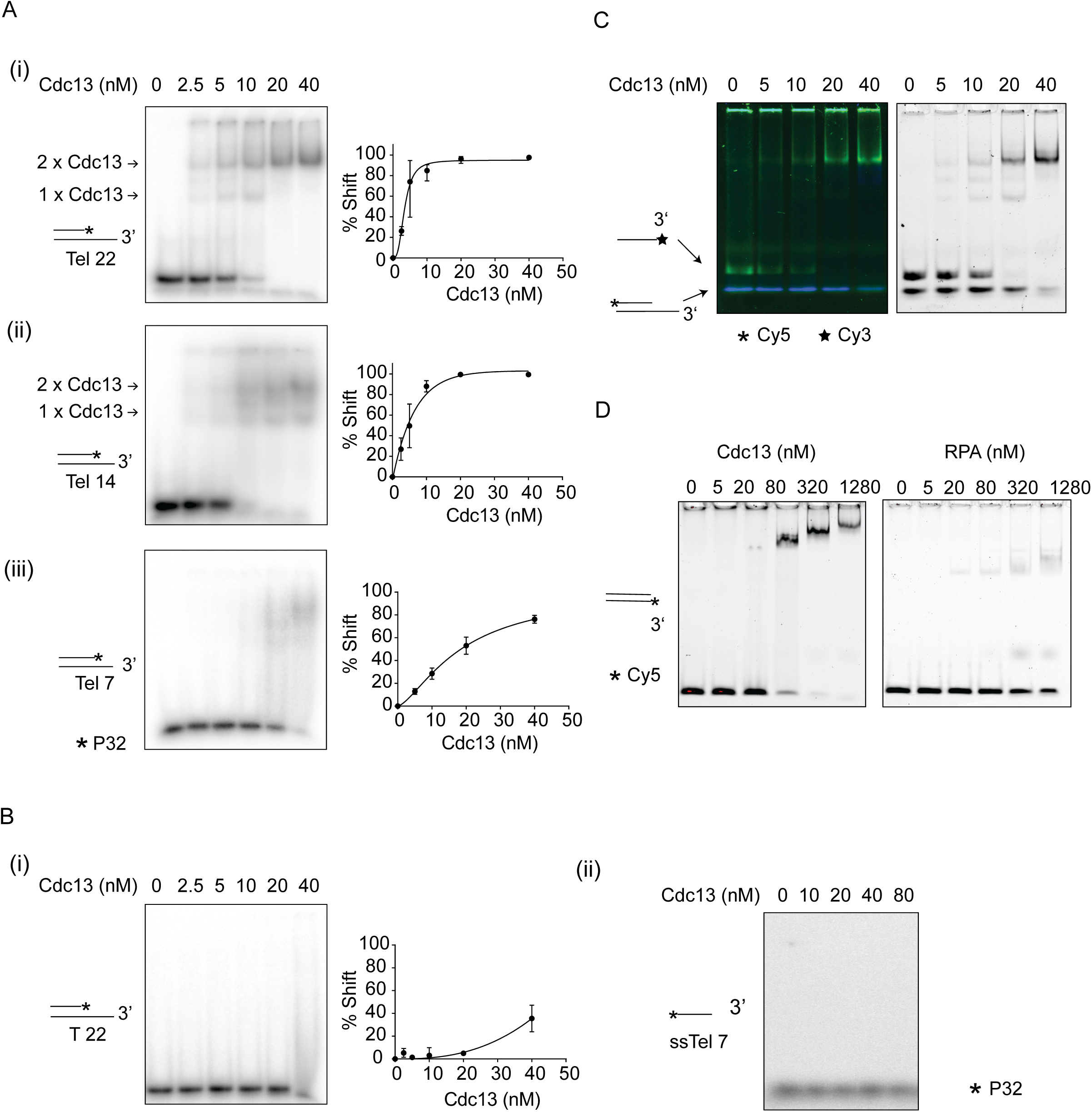
Cdc13 preferentially binds telomeric 3′ overhangs and ss/dsDNA junctions. (A) EMSA showing binding of Cdc13 to 5′- ^32^P-labeled telomeric substrates: Tel22 (i), Tel14 (ii), and Tel7 (iii) (5 nM each). Mean values ±SD from three independent experiments were plotted. (B) EMSA showing binding of Cdc13 to 5′-^32^P-labeled T22 (i) and ssTel22 (ii) substrates (5 nM each). Mean values ±SD from three independent experiments were plotted. (C) Competitive EMSA in which 3’-Cy5-labeled Tel22 competes with 3’-Cy3-labeled ssTel22 or excess ssDNA. Mean values ±SD from three independent experiments were plotted. (D) EMSA comparing Cdc13 and RPA binding to a Cy5-labeled 26-bp blunt-end dsDNA (5 nM).

To confirm this, we conducted a competition assay of two substrates: ssDNA of telomeric sequences (ssTel22) and 3’ overhang of telomeric sequences (Tel22). When both substrates were present in EMSA, Cdc13 strongly favored Tel22 over ssTel22 (**Fig. 1C**), suggesting that junction structure contributes to overall affinity. We next directly assessed Cdc13 binding to a blunt-ended 26-bp duplex substrate (ds26) and observed that Cdc13 binds ds26 with an apparent Kd of ∼300 nM. This interaction is specific and was not observed with Replication Protein A (RPA) (**Fig. 1D**), the universal ssDNA binding protein in eukaryotes^41^, confirming Cdc13’s capacity to engage double-stranded DNA. In summary, these results demonstrate that Cdc13 binds telomeric 3′ overhangs with sequence and length dependence, and that duplex recognition at the ss/dsDNA junction enhances its binding affinity. We surmise that this dual engagement mode distinguishes Cdc13 from RPA and may underlie its function in telomere protection.

### Conserved lysine 504 is essential for telomeric ss/dsDNA junction recognition by Cdc13

To explore the structural basis of duplex recognition by Cdc13, we first examined available structural information of its DNA-binding domain. Analysis of the *K. lactis* Cdc13 OB2-OB4 crystal structure (PDB: 6LBR)^42^ revealed an auxiliary α-helix near the OB3 N-terminus. This helix positions its side chains toward the phosphate backbone of 5’-terminal nucleotides A1-A5 (**Fig. 2A(i)**). Notably, a lysine residue (K521) in this helix is positioned 3.3 Å from the DNA backbone, suggesting possible electrostatic interactions (**Fig. 2A(i)**). Sequence alignment confirmed that this lysine is evolutionarily conserved among fungal Cdc13 homologs (**Fig. S2A**), and the spatial arrangement is preserved in the *S. cerevisiae* Cdc13 OB3 structure (PDB: 1KF1^43^, **Fig. 2A(ii**)).

**Figure 2.**
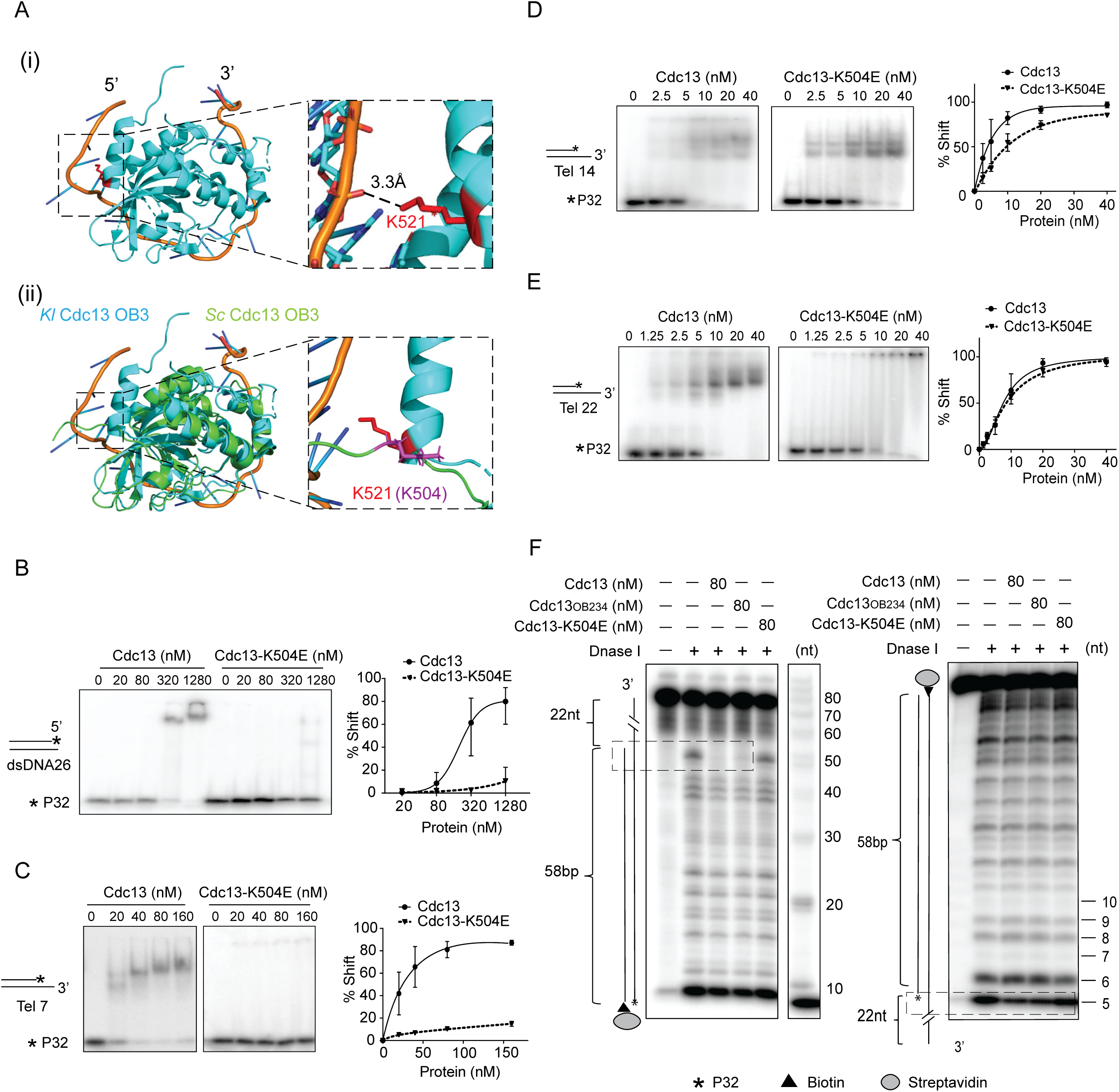
Lysine 504 is required for junction recognition by Cdc13. (A) Structural comparison of *K. lactis* and *S. cerevisiae* Cdc13 DNA-binding domains. (i) Crystal structure of the *K. lactis* Cdc13OB3–Tel25 complex (PDB: 6LBR) showing Lys521 (red) positioned 3.3 Å from the DNA phosphate backbone near the 5′ end of the duplex. (ii) Structural alignment of *K. lactis* (cyan) and *S. cerevisiae* (green) Cdc13 DNA binding domains (OB3), highlighting the conserved Lysine (K521 in *K. lactis* and K504 in *S. cerevisiae*). Structures were visualized and aligned using PyMOL. (B-E) EMSA comparing Cdc13 and Cdc13-K504E binding to a 5′- ^32^P-labeled 26-bp blunt-end dsDNA (B), Tel7 (C), Tel14 (D), Tel22 (E). Quantification curves represent mean ± SD from three independent replicates. (F) Localization of Cdc13, Cdc13OB234, and Cdc13-K504E at the ssDNA–dsDNA junction determined by DNase I footprinting assay. L-Tel22 was labeled with ^32^P on the blunt end. Protected regions are boxed; sizes indicated in nucleotide.

Based on the alignment, we hypothesized that the lysine K504 in *S. cerevisiae* Cdc13 may play a role in recognizing the ss/dsDNA junction upon binding to telomeric 3′ overhang.

To test this hypothesis, we generated and purified the K504E point mutant of full-length Cdc13 (**Fig. S2B**), which displayed a similar monomer/dimer distribution as the wild-type protein in size-exclusion chromatography (**Data not shown**). EMSAs using either the duplex DNA substrate ds26 or the minimal telomeric ss/dsDNA junction mimic Tel7 showed that Cdc13-K504E displayed minimal binding to both substrates compared to wild-type (**Fig. 2B** and **C**), consistent with impaired junction engagement. In contrast, binding to longer telomeric overhangs (Tel22 and Tel14) (**Fig. 2D and 2E**), as well as fully single-stranded ssTel22 substrate (**Fig. S2D**), was largely retained, suggesting that the mutation selectively disrupts duplex or junction recognition while preserving ssDNA binding.

### Telomeric ss/dsDNA junction recognition by Cdc13

To further examine the binding and protection of telomeric ss/dsDNA junctions by Cdc13, we performed DNase I footprinting using L-Tel22, a substrate consisting of a longer duplex region (58 bp) with a 22-nt 3′ telomeric overhang. Cdc13 protected approximately 3-5 bp of duplex DNA immediately at the ss/dsDNA junction (**Fig. 2F**), consistent with prior reports ^44^. Notably, the OB234 fragment of the *S. cerevisiae* Cdc13 (**Fig. S2**C) also retained duplex protection, suggesting OB1 is not involved in junction recognition. In contrast, the Cdc13-K504E mutant completely lost the duplex protection, indicating a failure to stably engage the duplex region at the junction. These findings affirm the critical role of K504 in Cdc13’s ability to recognize telomeric ss/dsDNA junctions^38^.

### Cdc13 suppresses Exo1-catalyzed DNA end resection by junction protection

As a crucial telomere end-protection factor, Cdc13 is required to safeguard mature telomeres against the Exo1 and Sgs1-Dna2 resection machinery in cells^13^. To determine how Cdc13’s junction binding contributes to this protection, we first assessed Cdc13’s impact on Exo1 nuclease activity using 3’-overhang substrates of various lengths and sequences. On Tel22 and Tel14 substrates, addition of 20 nM Cdc13 significantly inhibited the initial cleavage by Exo1 (3-fold inhibition, *p* < 0.0001, n = 3) (**Fig. 3A, S3A**). The effect was sequence specific as Cdc13 had no significant effect on Exo1 cleavage of T22 (*p* = 0.15, n = 3) (**Fig. 3B**). In addition, no direct interaction was detected between Cdc13 and Exo1 by pull-down (**Fig. S3B**), suggesting that Cdc13 exerts its function through DNA binding alone. Interestingly, RPA, which binds ssDNA but lacks junction specificity, only modestly affected Exo1 activity on tested substrates (<20% inhibition), supporting that the strong Cdc13 inhibitory effect on Exo1 is not only from its ssDNA binding (**Fig. 3A, S3A**). We previously reported that duplex recognition is essential for Exo1 activation^45^. Indeed, the K504E mutant, which is defective in duplex recognition, is largely defective in Exo1 inhibition on both Tel22 and Tel14 substrates (**Fig. 3C, S3C**). Hence, Cdc13’s junction binding may competitively exclude Exo1 from recognizing its substrate DNA, in a manner redundant with the Ku complex^46^.

**Figure 3.**
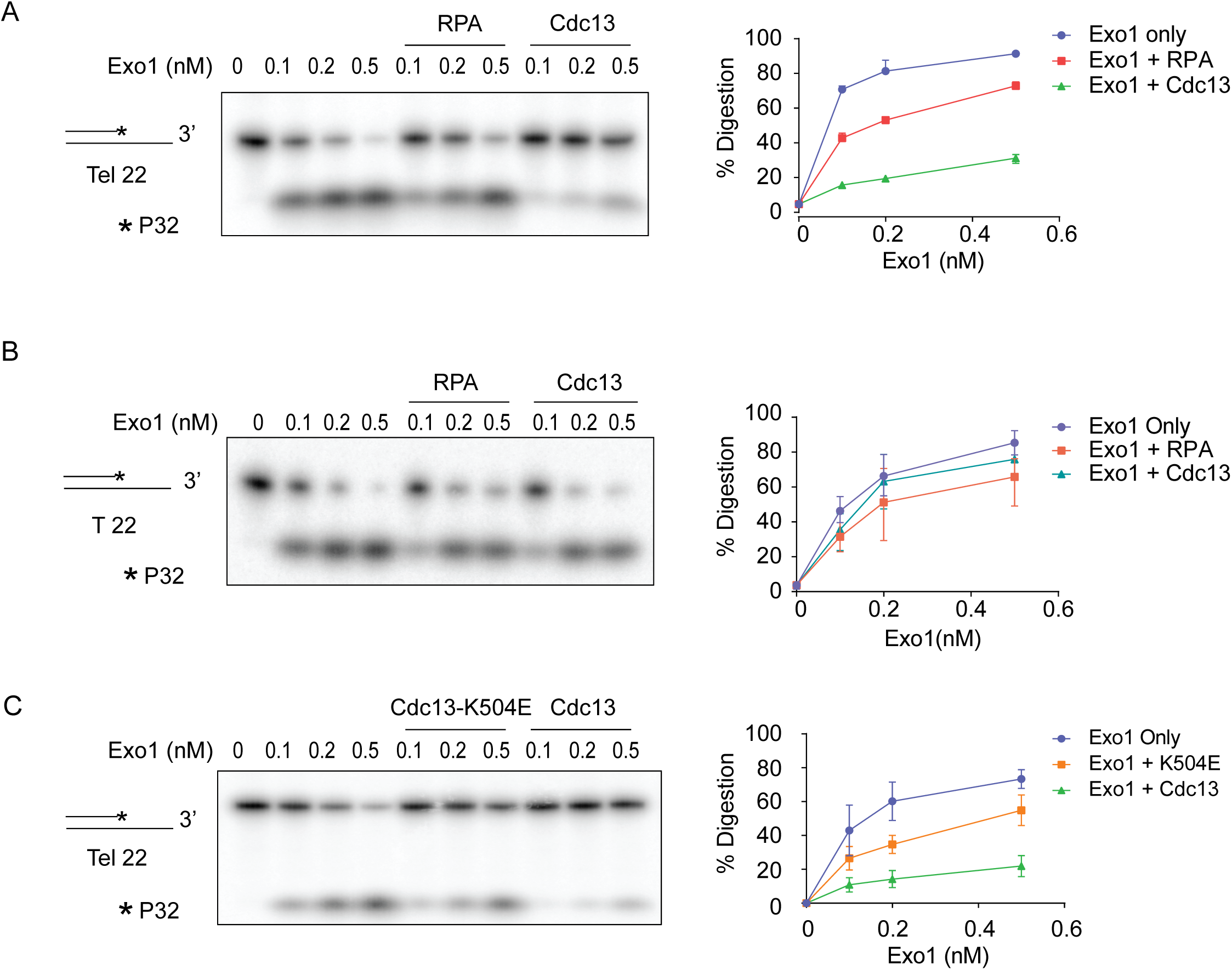
Cdc13 inhibits Exo1 via junction engagement. Exo1 resection assays using ³²P-labeled Tel22 (A, C) or non-telomeric T22 (B) substrates in the presence or absence of Cdc13, RPA, or Cdc13-K504E, as indicated. Reaction products were analyzed by denaturing PAGE and visualized by phosphorimaging. Quantification represents mean ± SD from three independent experiments.

### Cdc13 synergizes with Ku to suppress Sgs1-Dna2 catalyzed DNA end resection

Beyond Exo1-mediated resection, Sgs1 helicase and Dna2 nuclease constitute another key machinery for DNA end processing, where RPA serves as an essential component^47,48^. To understand how Cdc13 regulates this Sgs1-Dna2-RPA resection pathway, we first investigated the impact of Cdc13 on the activities of each component in this pathway. Using Tel7, Tel14, and Tel22 as substrates, we found that Sgs1 unwound the telomeric overhangs with a preference for a longer 3’ overhang (**Fig. 4A**). The addition of RPA substantially stimulated unwinding, showing the strongest enhancement on the Tel7 substrate (∼6-fold increase). On the other hand, Cdc13 (20 nM) mildly inhibited Sgs1-mediated unwinding, showing 2-fold reduction (**Fig. 4A(i)**).

**Figure 4.**
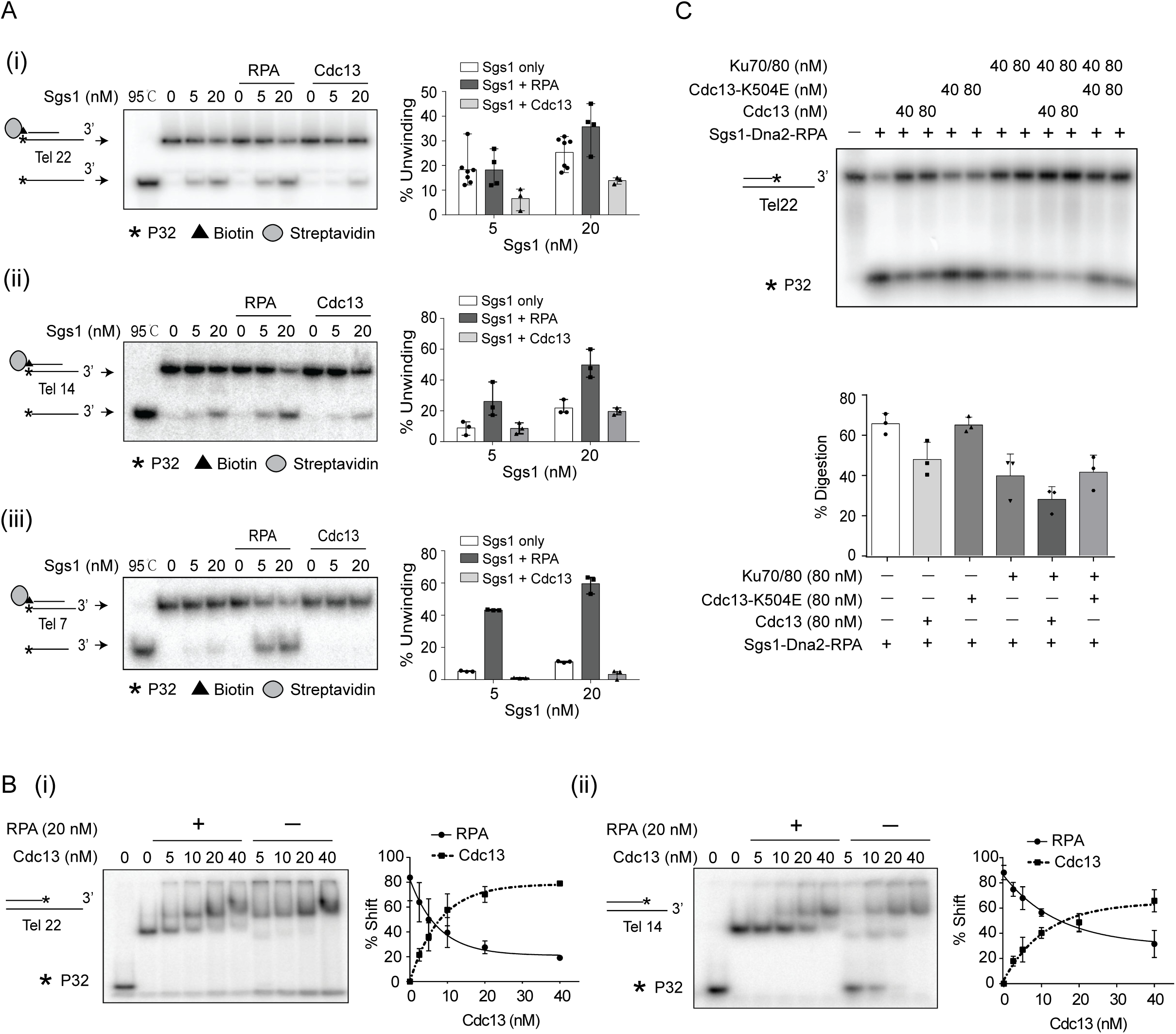
Cdc13 suppresses Sgs1-Dna2 catalyzed DNA end resection (A) ^32^P-labeled Tel22 (i), Tel14 (ii), and Tel7 (iii) substrates were incubated with Sgs1 in the presence of RPA or Cdc13 to assess the effect of each protein on Sgs1-mediated unwinding, followed by analysis on native TBE gels. Quantification represents mean ± SD from three independent experiments. (B) ^32^P-labeled telomeric substrates Tel22 (i) and Tel14 (ii) were first incubated with 20 nM RPA, followed by increasing concentrations of Cdc13. Reaction mixtures were analyzed on native TBE gels. Quantification represents mean ± SD from three independent experiments. (C) Resection reactions were assembled by first incubating ^32^P-labeled telomeric DNA substrates with Ku, followed by the addition of Cdc13 or Cdc13-K504E. Resection was initiated by adding Sgs1 (5 nM), Dna2 (5 nM), and RPA (20 nM). Reaction products were analyzed on denaturing PAGE. Quantification represents mean ± SD from three independent experiments.

Dna2 is a ssDNA endonuclease. RPA stimulates its 5’ cleavage but inhibits its 3’ cleavage^49,50^. To compare Cdc13 with RPA, we first asked how Cdc13 affects Dna2-catalyzed 5’ C-strand digestion which occurs during telomeric DNA end resection. To do this, we tested Dna2 nuclease activity without or with Cdc13 or RPA on a 5’-C-rich overhang (5’-C22) and included a 5’-G-rich overhang (5’-Tel22) as a control. While RPA stimulated cleavage of both substrates by Dna2, Cdc13 had little effect on 5’-C22 cleavage (**Fig. S4A(i)**) likely due to its low affinity for the C-strand ssDNA. In contrast, Cdc13 efficiently inhibited Dna2 activity on 5’-Tel22 (**Fig. S4A(ii)**), indicating Cdc13 is not a Dna2 functional pair like RPA. Next, we compared Cdc13 and RPA in the protection of telomeric 3’ overhang from Dna2 digestion. In this case, both RPA and Cdc13 protected the 3’ end of Tel22 (a 3’-G-rich overhang) from Dna2 digestion (**Fig. S4A(iii)**). These data suggest that while Cdc13 has little effect on Dna2-catalyzed 5’ C-strand digestion, it can substitute for RPA in the protection of nascent 3’ telomeric G-strand ssDNA.

Given RPA’s essential role in the Sgs1-Dna2 pathway and the opposing effects of Cdc13 versus RPA on Sgs1 and Dna2 activities, we asked how do Cdc13 and RPA compete in binding to telomeric DNA using EMSAs. Under physiological salt (150 mM KCl), Cdc13 effectively outcompeted RPA for Tel22/Tel14 binding (**Fig. 4B**), but, interestingly, only in its dimer form. This dominance aligns with Cdc13’s superior binding affinity for telomeric DNA. In summary, our experiments show that Cdc13 binds the 3’ G-strand and competes with RPA, which limits Sgs1- catalyzed unwinding of the 5’ end, thereby suppressing end resection. At the same time, occupying the 3’ strand by Cdc13 also shields the 3’ overhang from Dna2 cleavage, protecting telomere integrity.

Given the reported synergy between Sgs1, Dna2 and RPA^47,51,52^, we examined the effect of Cdc13 in a reconstituted DNA end resection reaction containing Sgs1, Dna2, and RPA with the Tel22 substrate. In this system, Cdc13 partially suppressed resection by Sgs1-Dna2-RPA, as predicted. Interestingly, the Ku complex, another telomere protection factor in yeast, synergized with Cdc13 in suppressing resection (**Fig. 4C**). This synergy requires sequential loading of Ku followed by Cdc13, to allow complex assembly at the telomeric DNA end (**Fig. S4B**) ^53^. Notably, the Cdc13- K504E mutant showed little effect either alone or in combination with Ku (**Fig. 4C**), indicating that junction-binding capacity is essential for Cdc13 to protect telomeric DNA ends against the Sgs1-Dna2 resection pathway.

### Cdc13 modulates Ku binding at telomeric ss/dsDNA junctions

The yeast Ku complex preferentially binds ss/dsDNA junctions *in vitro* ^46,54,55^, which could create a spatial conflict with Cdc13 on telomere. High-resolution ChIP at telomeres showed that Ku localizes to distal internal telomeric sequences (ITSs) further from chromosome ends^20^ , leading us to hypothesize that Cdc13 may displace Ku at the junction to prevent Ku mediated end joining. To test this hypothesis, we examined the Cdc13 and Ku occupancy at the ss/dsDNA junction of telomeric DNA using a DNase I footprinting assay (**Fig. 5A**). In this assay, Cdc13 efficiently protected 3-5 bp at the junction site, consistent with our observations above. The protection was abolished by the K504E mutation, again confirming the role of K504 in junction recognition. The Ku complex, on the other hand, protected ∼15 bp near the junction (**Fig. 5A**), consistent with a previously published result^46^. Interestingly, when Ku and Cdc13 were sequentially loaded, the protection at the vicinity of junction by Cdc13 remains, while the rest of the region protected by Ku was broadened and showed generally reduced protection, which may be caused by the displacement of Ku from the junction (**Fig. 5A**). Again, the signature of Ku displacement was lost when using the Cdc13-K504E mutant. Hence, this redistribution of the Ku protection pattern indicates that Cdc13 remodels the pre-bound Ku complex, displacing it from the ss/dsDNA junction, an effect that requires its junction binding capacity.

**Figure 5.**
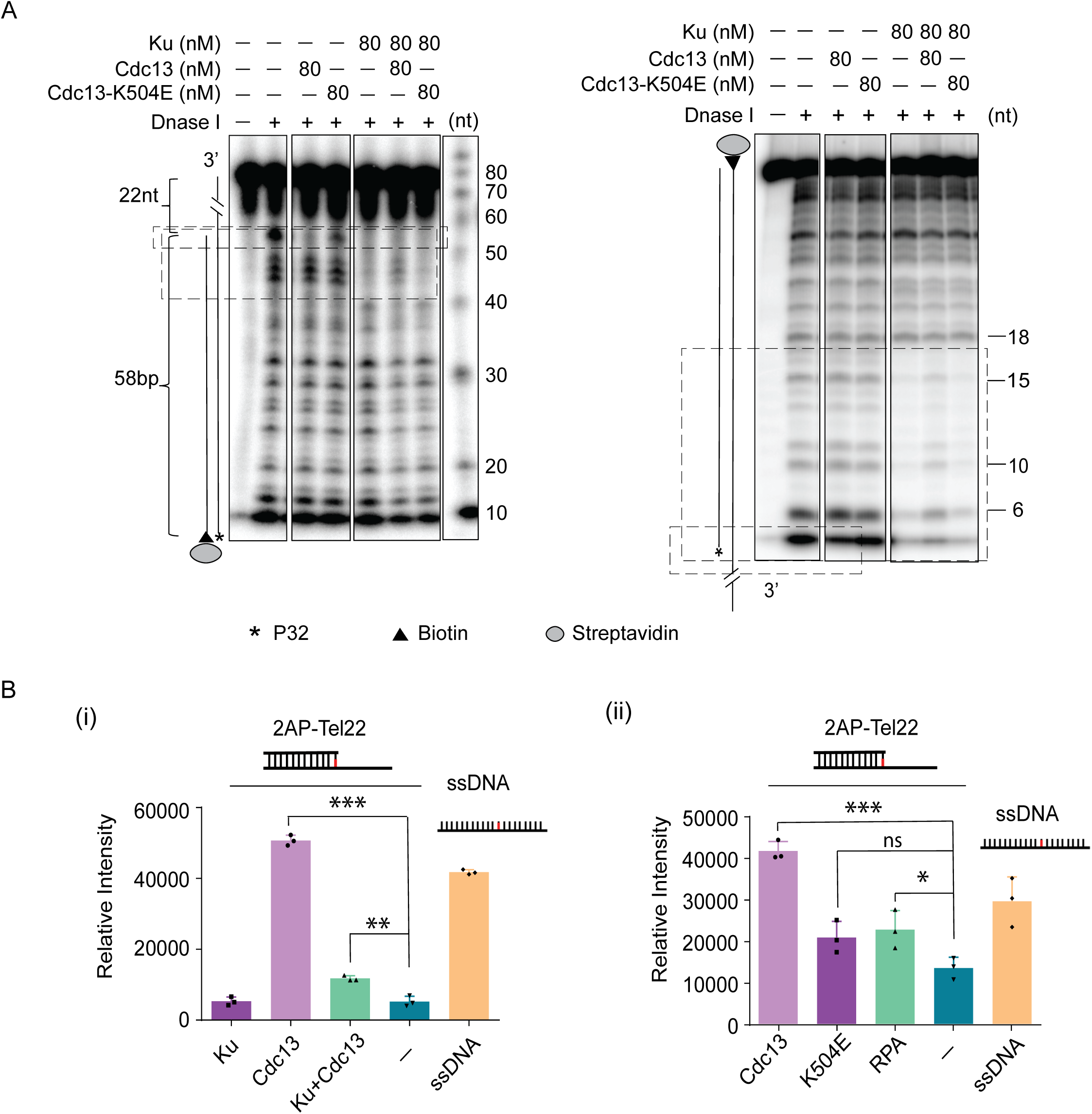
Cdc13 remodels Ku localization at the telomeric ss/dsDNA junction. (A) Localization of Cdc13 or Cdc13-K504E in the presence of Ku at the ssDNA–dsDNA junction determined by DNase I footprinting assay. L-Tel22 was ³²P-labeled at the blunt end. Protected regions are boxed, with sizes indicated in nucleotides. (B) Tel22 contains a 5′ nucleotide substituted with 2-aminopurine (2AP), an adenine analog that reports on base-pairing and base-fliping interactions. 2AP fluorescence increases upon Cdc13 binding, reaching a higher level compared with Ku (i), Cdc13-K504E (ii), or RPA (ii). The combination of Cdc13 and Ku further enhances the fluorescence relative to Ku alone (i).

To explore the mechanism underlying Ku repositioning, we employed a 2-aminopurine (2AP) base-flipping assay to monitor structural changes at the 5′ end of the junction. 2AP is an adenine analog that fluoresces weakly when base-stacked in duplex DNA but emits strongly (∼370 nm) upon base flipping, thus serving as a sensitive reporter of local DNA distortion^56,57^. When 2AP was placed at the first base pair of the junction duplex region, Cdc13 markedly increased 2AP fluorescence (*p* < 0.0001, n = 3), indicating local base flipping (**Fig. 5B(i)**). This signal was specific to Cdc13 and was not observed with Cdc13-K504E or RPA (p > 0.1 and p = 0.039, respectively, n=3). When Ku and Cdc13 were sequentially loaded, the enhanced fluorescence persisted, whereas Ku alone had no effect, suggesting that the conformational change is primarily driven by Cdc13 (p < 0.01, n = 3) (**Fig. 5B(ii)**).

Together, these results indicate that Cdc13 repositions Ku at the telomeric ss/dsDNA junction by remodeling the DNA end structure, a process that involves local base flipping.

### *cdc13-K504E* and Ku mutants are sensitive to resection stress

Having established that Cdc13-K504E impairs junction recognition and resection inhibition *in vitro*, we next asked whether these defects translate to physiological consequences in cells. Surprisingly, although *CDC13* is an essential gene, *cdc13-K504E* mutant is viable and not temperature sensitive, unlike certain other *cdc13* alleles such as *cdc13-1*(**Fig. S6A, S5B**) ^58,59^. Moreover, no significant telomere shortening was observed in the *cdc13-K504E* mutant **(Fig S5C).** Interestingly, the *cdc13-K504E ku80Δ* remains viable but displays a temperature-sensitive phenotype and telomere shortening similar to that observed in *ku80Δ* (**Fig. S5A, S5B, S5C**) ^17,60,61^, indicating the plasticity of telomere end protection in yeast. To further challenge telomeric DNA protection, we overexpressed Exo1 using a galactose inducible system. While overexpression of Exo1 resulted in a mild growth defect in *cdc13-K504E* and *ku80Δ* single mutants (**Fig. 6A**), the *cdc13-*K504E *ku80Δ* double mutant showed a synthetic growth defect upon Exo1 overexpression, with viability reduced by over 100-fold compared to the wild type (**Fig. 6A**). The findings suggest that Cdc13 and Ku play redundant roles in suppressing Exo1-mediated resection in cells.

**Figure 6.**
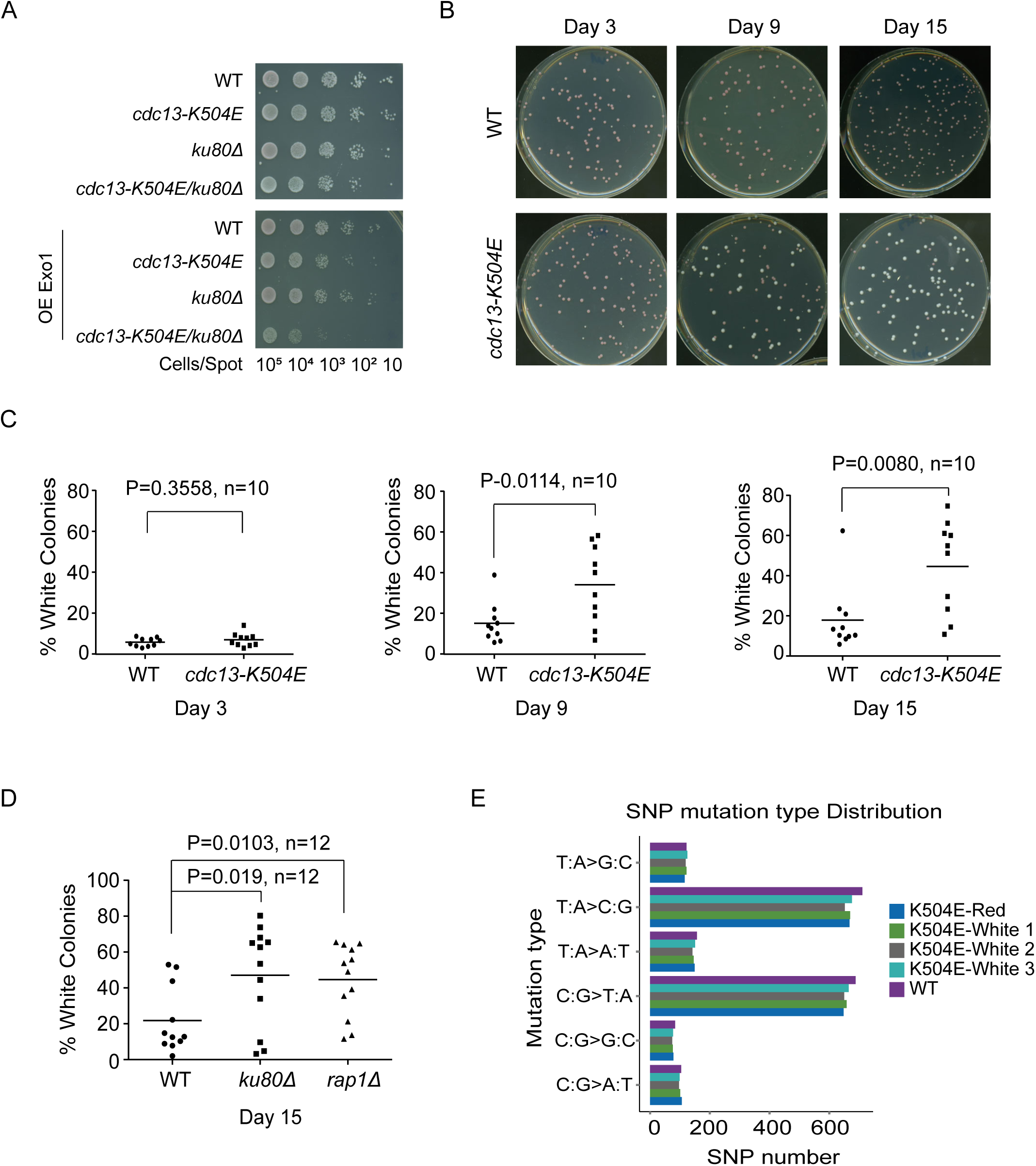
*cdc13*-K504E generates adaptive *ADE2* revertants during prolonged stationary phase (A) Growth assay of wild-type, *cdc13*-K504E, *ku80Δ*, and *cdc13*-K504E *ku80Δ* strains with Exo1 overexpression from plasmids. Cells were spotted onto selective raffinose or galactose plates at 30°C. (B) Colony color reversion assay. wild-type and *cdc13*-K504E strains were cultured in liquid YPD for 3, 9, or 15 days, then plated on fresh YPD plates. Representative images are shown. (C) Quantification of white colony frequency for wild-type and *cdc13*-K504E strains from 10 independent YPD plates on days 3, 9, and 15. Each point represents an independent plate; horizontal bars denote the mean. Statistical significance (*P* values) is indicated. (D) Quantification of white colony frequency for wild-type, *ku80Δ* and *rap1Δ* strains from 10 independent YPD plates on day 15. Each point represents an independent plate; horizontal bars denote the mean. Statistical significance (*P* values) is indicated. (E) Genome-wide distribution of single-nucleotide polymorphism (SNP) mutation types. Shown are wild-type and four *cdc13-*K504E isolates (three white colonies and one red colony) identified by whole-genome sequencing.

### *cdc13-K504E* mutant is prone to the formation of *ADE2* revertants with a fitness gain during stationary phase growth

To assess the long-term consequences of impaired telomere capping in the *cdc13-K504E* mutant, we subjected wild-type and *cdc13-K504E* cultures to prolonged stationary phase culturing (15 days in YPD without dilution). We first evaluated cellular resilience by plating 200 cells per replicate after chronic stress exposure. Both strains exhibited comparable colony-forming efficiency after 15 days (**Fig. S6D**), indicating no gross defect in recovery capacity in the *cdc13-K504E* mutant.

Unexpectedly, during this analysis, however, we observed an interesting phenomenon: *cdc13- K504E* plates showed progressive accumulation of white, non-pigmented colonies (**Fig. 6B**), whereas WT controls remained predominantly red. Quantitative analysis revealed that white colonies increased progressively with extended incubation, reaching approximately 35% by day 9 and 45% by day 15, whereas wild-type controls remained largely red, with white colonies comprising no more than 18% of the population under the same conditions (**Fig. 6C**). Deletion of *RAP1* or *KU80*, two other telomere-protection factors, also led to a marked increase in white colony formation (**Fig. 6D**), supporting the association of this phenotype with telomere dysfunction.

In the yeast strain background used in this study (W303), colonies appear red due to the *ade2-1* nonsense mutation, which disrupts adenine biosynthesis and leads to pigment accumulation^62,63^. Restoration of *ADE2* function reverts colony color to white. To confirm the genetic basis of this pigmentation switch, we sequenced the *ADE2* locus from eight independent *cdc13-K504E* white colonies. All eight harbored reversion mutations restoring the *ADE2* open reading frame (**Fig. S5E**).

The accumulation of *ADE2* revertants during continuous stationary phase culturing could be due to either a mutator phenotype or a fitness gain of these *ADE2* revertants. The latter possibility seems more likely, as we consistently observed an increase in the final cell density of the culture with *ADE2* revertant enrichment. Furthermore, whole-genome sequencing (WGS) of wild-type red (WT-red), *cdc13-K504E* red (K504E-red), and *cdc13-K504E*-derived white colonies (K504E- white) revealed comparable numbers and spectra of SNPs across samples (**Fig. 6E**), ruling out the mutator phenotype in the *ADE2* revertant.

### Fitness gain in the *ADE2* revertants is inheritable and likely associated with transcriptomic reprogramming

In addition to the fitness gain during stationary phase growth, we noticed a larger colony size of the majority of *ADE2* revertants after plating (**Fig. S5F**). Growth curve analysis showed white colonies grew faster and to higher densities than red ones (**Fig. S6A**), indicating the inheritance of the fitness gain under normal vegetative growth. Fluorescence-activated cell sorting (FACS) of asynchronous cultures revealed that K504E-white cells exhibited a decrease in the G1 population from ∼35% to ∼30% and a corresponding increase in G2 from ∼40% to ∼45%, while the S-phase fraction remained largely unchanged **(Fig. S6B)**, consistent with faster G1 progression.

Despite the inheritance of the fitness gain, to our surprise, whole-genome sequencing of three white clones didn’t reveal consensus mutations, except for the *cdc13-K504E* mutation and the *ADE2* reversion. To uncover the molecular basis of the fitness gain, we compared the transcriptomes of wild-type red (WT-red), *cdc13*-K504E red (K504E-red), and *cdc13*-K504E white (K504E-white) colonies by RNA sequencing. Principal component analysis showed clear segregation among these groups (**Fig. S6C**), confirming distinct transcriptional states. Comparisons of differentially expressed genes revealed 975 genes commonly affected by the K504E mutation, while 270 genes differed between red and white colonies, confirming progressive transcriptomic divergence (**Fig. S6D**). Among them, there is a broad upregulation of metabolic genes in K504E-white colonies, encompassing critical pathways in adenine biosynthesis (e.g., *ADE1, ADE2, ADE13, ADE17 and ADE57*) and thiamine biosynthesis/salvage/intake (e.g., *THI4*, *THI20*, *THI21, THI73 and PET18*) (**Fig. 7A, B**). Genes showing significant transcription differences were further validated by RT–qPCR of key targets (**Fig. S6E**). Notably, no detectable mutations were found within the differentially expressed genes or their flanking regions in our whole-genome sequencing data, suggesting that the observed expression changes are due to transcriptional regulation rather than local genetic alterations.

**Figure 7.**
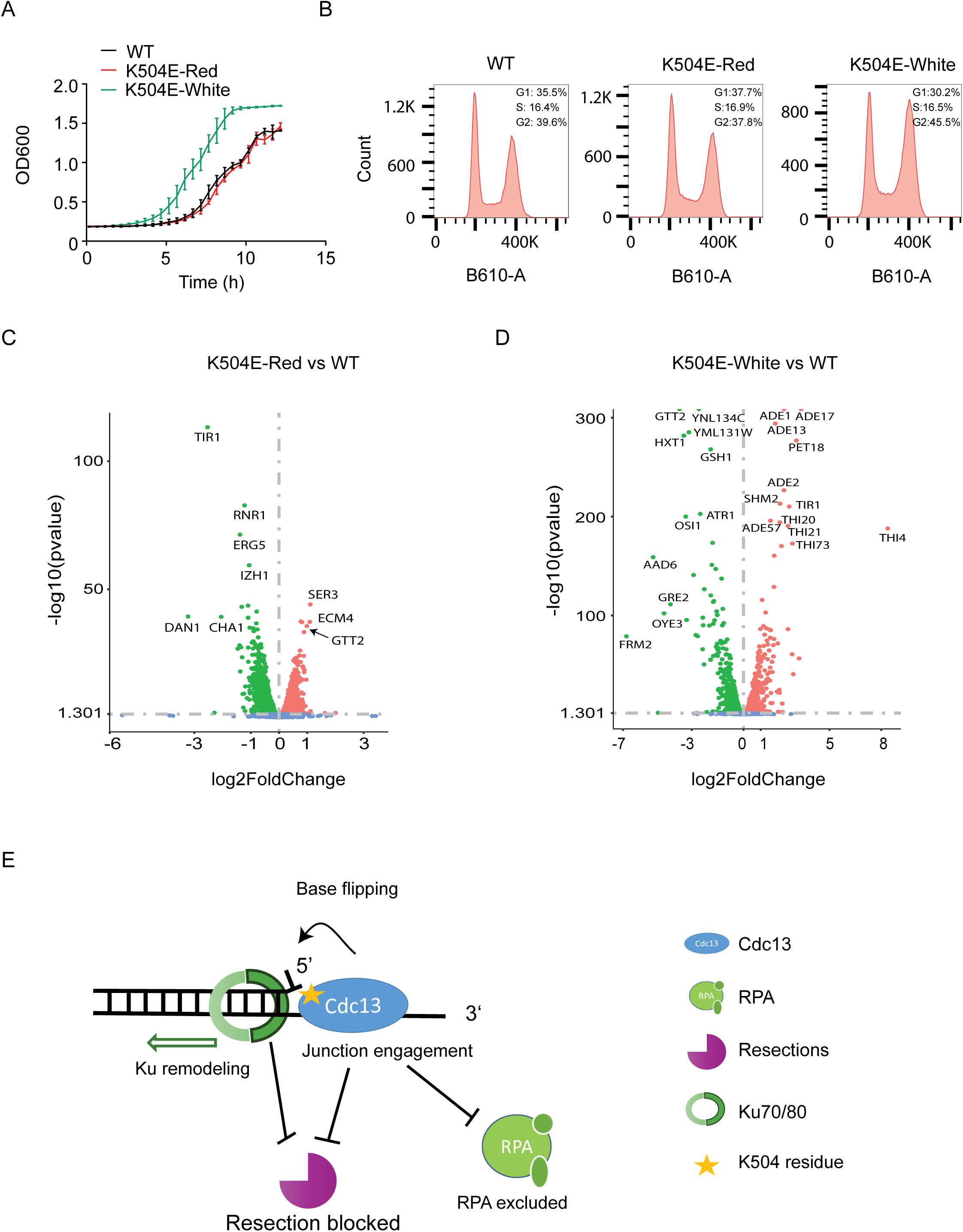
White *cdc13*-K504E colonies exhibit altered cell cycle progression and DNA content profiles. (A-B) Volcano plot comparing K504E-red vs WT (A) and K504E-white vs WT (B). (C) Schematic model illustrating proposed Cdc13–Ku interaction at the telomeric ss/dsDNA junction.

In yeast, thiamine (vitamin B1) acts as an essential cofactor, primarily as thiamine pyrophosphate (TPP), for enzymes involved in glycolysis, the pentose phosphate pathway, and redox homeostasis^64,65^. Notably, *THI4*, the most upregulated gene identified in our RNA-seq analysis, encodes thiazole synthase, which catalyzes a rate-limiting step in thiamine biosynthesis^66^. Previous studies have shown that *THI4* overexpression promotes faster cell growth and enhanced fitness, likely by accelerating glucose uptake, boosting glycolytic flux, and increasing stress tolerance^67^.

## Discussion

Our study characterized a dual DNA-binding activity of the yeast telomeric protein Cdc13 that underlies its ability to inhibit nucleolytic degradation and inappropriate DNA repair. We found that in addition to its established high-affinity binding to the single-stranded telomeric 3′ overhang, Cdc13 also interacts with the adjacent duplex DNA at the ss/dsDNA junction via a conserved lysine residue (K504). This junction engagement is critical for suppressing Exo1- and Sgs1–Dna2- mediated resection and safeguarding chromosome ends.

Despite this defect, *cdc13-K504E* mutants do not exhibit immediate telomere shortening. This apparent resilience likely reflects functional buffering by other telomere-associated factors such as Rap1-Rif1/2 and the Ku complex, which also help maintain telomere structure and delay overt genome instability^4^. The delayed emergence of phenotypes suggests that junction-binding may contribute to long-term end protection, particularly under stress.

Although buffered in early passages, *cdc13-K504E* mutants ultimately exhibit cellular changes during prolonged stationary phase. White colonies derived from long-term culture show distinct transcriptomic reprogramming, with upregulation of thiamine and adenine biosynthetic pathways. Flow cytometry further revealed an enrichment of cells in G2 phases and reduced G1 population, consistent with faster cycling. Whole-genome sequencing showed no mutations near key upregulated genes, suggesting that the expression changes reflect regulatory adaptation rather than local mutations. These observations raise the intriguing possibility that telomere erosion may serve as a timer to activate metabolic rewiring as an adaptive stress response. The *cdc13-K504E*, and other telomere protection mutants are more prone to this activation. This adaptive metabolic reprogramming echoes the link between telomere dysfunction and metabolic reprogramming observed in human cells^68–70^, offering insight into how telomere-driven metabolic changes contribute to cancer development, where metabolic plasticity is a hallmark of cancer^71^.

Biochemically, junction binding by Cdc13 repositions Ku localization at chromosome ends. Our footprinting and base-flipping assays suggest that Cdc13 reshapes duplex DNA near the junction, releases Ku complex from ss/dsDNA and promotes its diffusion along duplex DNA. This spatial reorganization, encompassing Ku displacement, RPA exclusion, and resection blockade, collectively safeguards telomere integrity (Fig 7C). Specifically, this reorganization achieves two critical outcomes: (i) synergistic enhancement of resection inhibition, and (ii) displacement of Ku from the ss/dsDNA junction. This displacement of Ku, while allowing it in resection protection, may also limit its capacity to catalyze NHEJ, thereby safeguarding telomere integrity by blocking both degradation and inappropriate repair.

This mechanism highlights a conserved protective strategy centered on ss/dsDNA junction recognition. However, its implementation differs between organisms. In budding yeast, which lacks a canonical shelterin complex, the CST component Cdc13 acts as a multi-functional hub, directly binding the ssDNA overhang, engaging the junction, inhibiting resection, and modulating Ku positioning. In mammals, both POT1 and the CST complex contribute to the inhibition of telomeric DNA resection^2,72,73^. Notably, POT1 exhibits ss/dsDNA junction recognition, analogous to yeast Cdc13’s junction-binding activity^73,74^, suggesting that while the division of labor among telomeric complexes has diverged, the ability to protect DNA ends at ss/dsDNA junctions is evolutionarily conserved. Our detailed mechanistic dissection of how Cdc13 safeguards the junction to prevent degradation and constrain aberrant repair in yeast therefore provides a model system for understanding how disruption of telomere-end integrity, whether during replication or end-protection, underlies the genomic instability characteristic of these human diseases.

## Materials and Methods

**Yeast Strains.** Yeast strains used in this study are derivatives of HKY3415-11 (*MATa leu2-3, 112 his3-11, 15 ade2-1, ura3-1, trp1-1, can1-100, RAD5+, SRS2-TAP-TRP1*) (A kind gift from Dr. Hannah Klein, NYU). All strains are listed in Supplementary Table S2.

**Yeast Growth Assays.** Yeast strains (Table S2) were cultured in YPD medium (1% yeast extract, 2% peptone, 2% dextrose) at 30°C with shaking. Strains were maintained on YPD or -LEU plates as required.

For spot assays, overnight saturated yeast cultures were harvested and adjusted to an OD600 of 1. Cells were then serially diluted 10-fold and spotted onto appropriate plates. The plates were air-dried and incubated at 30°C.

For growth curve analysis, overnight cultures were diluted to OD600 = 0.2 and grown to the exponential phase. Exponentially growing cells were further diluted to OD600 = 0.2 in fresh medium and seeded into 96-well plates. OD600 was recorded every 30 minutes at 30℃ using a plate reader.

For white colony assays, single colonies were inoculated into 3 mL YPD and grown at 30°C with shaking (160 rpm, Excella E24 incubator shaker) for 15 days. Cultures were then plated onto fresh YPD plates and incubated at 30°C for 2 days, followed by storage at 4°C until colonies developed pigmentation. Colonies were visually inspected, and the proportion of white colonies was determined from at least 10 independent clones per genotype.

**DNA Substrates Preparation.** All oligonucleotides were purchased from IDT, with sequences listed in Supplementary Table S1. For 5′-end labeling, oligonucleotides were radiolabeled with [γ- ^32^P] ATP using T4 Polynucleotide Kinase (New England Biolabs). For 3′-end labeling, oligonucleotides were labeled with Cy5-dUTP or Cy3-dUTP using terminal transferase (New England Biolabs). Excess ATP/dUTP nucleotides were removed using Bio-Spin 6 columns (Bio-Rad) before annealing complementary oligonucleotides in NEB Buffer 3.1 (50 mM Tris-HCl, pH 7.9, 100 mM NaCl, 10 mM MgCl2, and 100 µg/mL BSA) following a standard thermal cycler protocol. Notably, the ssTel7 substrate was not subjected to Bio-Spin 6 cleanup due to its short length.

**Cdc13, Cdc13-K504E and Cdc13-OB234 Expression and Purification.** The pET28a–His-SUMO–Cdc13-Flag/pET28a–His-SUMO–Cdc13-K504E-Flag/pET28a–His-SUMO–Cdc13-OB234-Flag (341-924) construct was transformed into *E. coli* BL21 (DE3) codon plus cells. Transformed cells were plated on Luria-Bertani (LB) agar containing 50 µg/mL kanamycin and incubated overnight at 37°C. A single colony was picked and inoculated into 2 L of LB liquid medium supplemented with 100 µg/mL kanamycin and grown statically at 37°C overnight. The culture was then shaken at 150 rpm until the optical density at 600 nm (OD600) reached 0.6–0.8. Protein expression was induced by adding isopropyl-β-D-1-thiogalactopyranoside (IPTG) to a final concentration of 0.5 mM, followed by incubation at 18°C overnight. Cells were harvested by centrifugation at 5,000g for 20 minutes, and the resulting cell pellets were stored at −80°C.

For protein purification, cell pellets from 12 L of culture were thawed and resuspended in 250 mL of buffer (40 mM KH2PO4, 500 mM KCl, 10% glycerol, 0.01% NP-40, 1 mM TCEP, pH 7.4) supplemented with 1 mM PMSF and a protease inhibitor cocktail (5 µg/mL each of aprotinin, chymostatin, leupeptin, and pepstatin A). Cells were lysed on ice using sonication for 10 minutes, and the lysate was clarified by centrifugation at 20,000g for 20 minutes. The supernatant was applied to a 2 mL Ni-NTA resin column (GE Healthcare) equilibrated with K buffer (40 mM KH2PO4, 500 mM KCl, 0.1% NP-40, 1 mM TCEP, and 15 mM imidazole). The column was washed twice with 20 mL of K buffer (40 mM KH2PO4, 500 mM KCl, 0.01% NP-40, 1 mM TCEP, and 15 mM imidazole), and the protein was eluted with 15 mL of K buffer containing 200 mM imidazole. The eluate was diluted 1:1 with K buffer (0.01% NP-40, 1 mM TCEP) and incubated overnight at 4°C with 0.5 mL anti-Flag beads and 5 µg Upl1 protease for SUMO cleavage. The Flag beads were washed twice with 10 mL K buffer (40 mM KH2PO4, 500 mM KCl, 0.1% NP-40, and 1 mM TCEP) and then with 20 mL K buffer (40 mM KH2PO4, 500 mM KCl, 0.01% NP-40, 1 mM TCEP). Bound protein was eluted with 5 mL of K buffer containing 200 µg/mL FLAG peptide and concentrated to 1 mL using an Amicon Ultra centrifugal filter (30 kDa cutoff, Millipore). The concentrated protein was further purified by size-exclusion chromatography using a Superdex 200 Increase 10/300 GL column (GE Healthcare). Fractions containing the target protein were pooled, concentrated to 0.5–1 mg/mL, and aliquoted in 10 µL volumes. The aliquots were flash-frozen in liquid nitrogen and stored at −80°C for future use.

### Exo1 Expression and Purification

The pET21a-Exo1 plasmid^45^ was transformed into Rosetta (DE3) pLys cells. An overnight culture was diluted 1:50 into 12 liters of Luria-Bertani (LB) medium containing 100 µg/mL ampicillin. Protein expression was induced with 0.5 mM isopropyl 1-thio-β-d-galactopyranoside (IPTG) when the culture reached an OD600 of 0.6–0.8, followed by incubation overnight at 16 °C. Cells were harvested by centrifugation, yielding approximately 30 g of cell pellet, which was stored at -80 °C for future use.

All purification steps were conducted at 4 °C. The cell pellet was resuspended in 150 mL of lysis buffer (pH 7.4) containing 40 mM KH2PO4 and 150 mM KCl, 0.5 mM EDTA, 10% glycerol, 40 mM KH2PO4, 0.01% NP-40, 1 mM β-mercaptoethanol, and a cocktail of protease inhibitors. Cells were lysed by sonication, and the lysate was clarified by centrifugation at 20,000 × g for 20 minutes. The supernatant was then loaded onto a 5-mL sulfopropyl-sepharose column and washed with 50 mL of washing buffer (pH 7.4) containing 40 mM KH2PO4 and 150 mM KCl before Exo1 was eluted with 50 mL of elution buffer (pH 7.4) containing 40 mM KH2PO4 and 500 mM KCl.

The eluate was incubated with 400 µL of nickel-nitrilotriacetic acid resin for 1.5 hours, followed by washing the resin with 30 mL of wash buffer (pH 7.4) containing 40 mM KH2PO4, 500 mM KCl and 15 mM imidazole. Exo1 was eluted in three fractions of 0.4 mL each using a stepped imidazole gradient of 50, 100, and 200 mM. The purified Exo1 protein, which peaked in the fraction containing 200 mM imidazole, was aliquoted and stored at -80 °C for future experiments.

### Ku Expression and Purification

The experimental procedure closely followed the method described by Krasner et al. (2015)^46^. Protein expression was carried out in the protease-deficient yeast strain BJ5464. Purification was performed using a two-step approach: DEAE-Sepharose chromatography followed by FLAG affinity purification.

### RPA Expression and Purification

RPA was expressed and purified as described previously^50^.

### Dna2 Expression and Purification

Dna2 was expressed and purified as described previously^50^.

### Electrophoretic Mobility Shift Assays (EMSA)

Electrophoretic mobility shift assays were performed in a 10 μL reaction system containing 20 mM Tris-HCl (pH 8.0), 2 mM MgCl2, 150 mM KCl, 1 mM DTT, 200 μg/mL BSA, 5 nM DNA substrate, and the indicated amounts of proteins. The samples were incubated at 30°C for 10 minutes. For each reaction, 2 μL of 6x loading dye (10 mM Tris-HCl (pH 8.0), 0.025% (w/v) orange G, 50% glycerol) was added. For the competition assay between RPA and Cdc13/Cdc13-K504E, RPA was added first and incubated at 30°C for 10 minutes, followed by the addition of Cdc13 or Cdc13-K504E and an additional incubation at 30°C for 10 minutes. In the competition assay between ssDNA and the 3’ overhang, 5 nM of each substrate was pre-mixed, and Cdc13 was added, followed by a 10-minute incubation at 30°C. The samples were then fractionated on a 6% native polyacrylamide gel run in 0.5x TBE (Tris-borate-EDTA) buffer, followed by phosphor-imaging analysis.

### Exo1 Resection Assays

Exo1 nuclease assays were performed in a 10 μL reaction system containing 5 nM DNA substrate. The DNA substrate was first incubated with the indicated amounts of ssDNA binding proteins in the reaction buffer (20 mM Tris-HCl pH 8.0, 2 mM MgCl2, 150 mM KCl, 1 mM DTT, and 200 μg/mL BSA). The mixture was incubated at 30°C for 10 minutes before adding Exo1. The reaction was then incubated at 30°C for an additional 10 minutes. To terminate and deproteinize the reaction, 0.2% SDS and 0.5 mg/mL proteinase K were added, and the sample was incubated at 37°C for 5 minutes, followed by heat treatment at 95°C for 5 minutes in loading buffer (95% formamide, 0.025% (w/v) bromophenol blue, 0.025% (w/v) orange G, and 5 mM EDTA). The sample was then chilled on ice for 5 minutes and fractionated by 20% polyacrylamide denaturing gel electrophoresis in 0.5x TBE (Tris-borate-EDTA) buffer, followed by phosphor- imaging analysis.

### Sgs1 Unwinding Assays

DNA substrate (5 nM) was incubated with Sgs1 and the indicated proteins in a 10 μL reaction buffer containing 20 mM Tris-HCl (pH 8.0), 2 mM MgCl2, 100 mM KCl, 1 mM ATP, 1 mM DTT, and 200 μg/mL BSA at 30°C for 5 min. Reactions were stopped and deproteinized by the addition of 0.2% SDS, 0.5 mg/mL proteinase K, and 1 μM cold DNA, followed by incubation at 37°C for 5 min.

### Dna2 Nuclease Assays

Assays were performed as previously described (Niu et al., 2010)^52^, with minor modifications.

### Sgs1-Dna2 Resection Assays

Assays were conducted in a 10 μL reaction system containing 5 nM DNA substrate and the indicated amounts of DNA binding proteins, which were incubated at 30°C for 10 minutes the reaction buffer (20 mM Tris-HCl pH 8.0, 2 mM MgCl2, 100 mM KCl, 1 mM ATP, 1 mM DTT, and 200 μg/mL BSA) before adding the enzyme mixture. The enzyme mixture was pre-mixed to achieve final concentrations of 5 nM Sgs1, 5 nM Dna2, and 20 nM RPA in the reaction. The reaction was then incubated at 30°C for 10 minutes. To stop and deproteinize the reaction, 0.2% SDS and 0.5 mg/mL proteinase K were added, and the sample was incubated at 37°C for 5 minutes, followed by heat treatment at 95°C for 5 minutes in loading buffer (95% formamide, 0.025% (w/v) bromophenol blue, 0.025% (w/v) orange G, and 5 mM EDTA). The sample was subsequently chilled on ice for 5 minutes and fractionated by 20% polyacrylamide denaturing gel electrophoresis in 0.5x TBE (Tris-borate-EDTA) buffer, followed by phosphor- imaging analysis.

### DNase I Footprinting

The reaction (20 μL) was prepared in a buffer containing 10 mM Tris-HCl (pH 7.5), 2.5 mM MgCl2, and 0.5 mM CaCl2. Ku was first incubated with the DNA substrate for 10 minutes at 30°C, followed by the addition of Cdc13 or its mutants, and incubated again at 30°C. DNase I (0.05 U) was then added, and the reaction was further incubated for 5 minutes at 25°C. The reaction was terminated by boiling in formamide loading dye, followed by analysis on 15% a denaturing TBE polyacrylamide gel.

### *In Vitro* Competition DNA Binding Assay

Tel22 and Tel14 (5 nM each) were incubated with RPA (20 nM) in a reaction buffer containing 20 mM Tris-HCl (pH 8.0), 2 mM MgCl₂, 150 mM KCl, 1 mM DTT, and 200 μg/mL BSA. RPA was first incubated at 30°C for 10 minutes, followed by the addition of Cdc13 at varying concentrations (2.5, 5, 10, 20, and 40 nM) and further incubation at 30°C for another 10 minutes. Correlation curves and quantifications, including the mean and standard deviation, were derived from three independent experiments. Solid and dashed lines in the correlation curves represent Cdc13 and RPA, respectively.

### Telomere Southern Blot Analysis

Genomic DNA was extracted from Saccharomyces cerevisiae and digested overnight with XhoI. Digested DNA was separated on a 0.8% agarose gel and transferred onto a positively charged nylon membrane (Hybond-N+, Amersham) by capillary blotting in 10× SSC. DNA was UV crosslinked to the (120 mJ/cm²).

A telomeric probe (5’-ACACCCACACCCACACCCACACCC-3’) was labeled using the DIG High Prime DNA Labeling Kit (Roche). The probe was hybridized to the membrane in prewarmed hybridization buffer at 55°C overnight. The membrane was washed under stringent conditions and incubated with anti-DIG-AP antibodies (Roche). Signal detection was performed using chemiluminescent substrates, and images were acquired using an imaging system.

### 2-AP Fluorescent Assay

2AP fluorescent samples were excited at 315 nm, and emission was detected at 370 nm with a Photon-counting spectrometer PC1 (ISS Inc). The reaction buffer contained 20 mM Tris-HCl (pH 8.0), 2 mM MgCl2, 150 mM KCl, 1 mM DTT, and 200 μg/mL BSA. Experiments were performed under room temperature with Ku (80 nM) or/and Cdc13/Cdc13-K504E (80 nM)

### Exo1 and Cdc13 Affinity Pull-down Assay

Exo1-His6 (1.2 μg) and Cdc13-Flag (1.5 μg) were incubated in 50 μL binding buffer (K150, 1 mM DTT) at 4°C for 1.5 hours. After incubation, 10 μL of the reaction mixture was taken as the input sample. The reaction was then incubated with Flag-beads, followed by centrifugation at 10,000g for 5 minutes to collect the flow-through. Beads were washed three times with 0.8 ml wash buffer (K150, 1 mM DTT, 0.01% NP-40), and the wash fractions were collected. Bound proteins were eluted by incubating the beads in 1× SDS loading buffer at 95°C for 5 minutes.

### Flow Cytometry Analysis

Cells were cultured in YPD medium at 30°C with shaking until reaching a density of 0.5–1 × 10^7 cells/mL. Cells were pelleted and fixed in 70% ethanol at room temperature for 2 hours or overnight at 4°C. Fixed cells were washed twice with 50 mM sodium citrate buffer, then incubated in sodium citrate containing 0.1 mg/mL RNase A at 37°C for at least 1 hour. Subsequently, cells were stained with 20 μg/mL propidium iodide (PI) in sodium citrate buffer with RNase A, protected from light, at room temperature for 1 hour or overnight at 4°C. Before flow cytometry, cells were sonicated twice for 10 seconds at full power and diluted to approximately 5 × 10^5 cells/mL in sodium citrate buffer containing 5 μg/mL PI. Samples were then analyzed by flow cytometry to assess cell cycle distribution. Optional steps included microscopic verification of PI staining and proteinase K treatment (10 μL of 20 mg/mL at 55°C for 1 hour) if needed.

### RNA Extraction and Sequencing

Total RNA was extracted from yeast cell samples by Novogene Co., Ltd. (Beijing, China) using their standard protocols. The extracted RNA was quality-checked and used to prepare mRNA libraries. RNA sequencing was performed on an Illumina NovaSeq X Plus platform with paired-end 150 bp reads (PE150), generating at least 6 Gb of raw data per sample. Sequencing quality was assessed with Q30 scores ≥ 85%. Raw data processing, including quality control, alignment to the Saccharomyces cerevisiae W303 reference genome (downloaded from the Saccharomyces Genome Database on June 28, 2025), and gene expression quantification, was performed using Novogene’s internal pipelines.

### RT-qPCR

RT-qPCR was performed on a Bio-Rad CFX system using SYBR Green detection chemistry. Actin was used as the reference gene, and relative expression levels were calculated using the 2^-ΔΔCq method, with WT as the calibrator. For each condition, two technical replicates were run; values are reported as mean ± SD. Melt-curve analysis confirmed single products and NTC/NRT controls showed no amplification.

### RNA-seq Data Analysis

Normalized gene expression values were provided by Novogene. Raw and processed RNA-seq data have been deposited in the NCBI Gene Expression Omnibus (GEO) under accession GSE305992. Subsequent analyses, including calculation of gene distances to telomeres based on SGD genomic coordinates, and generation of heatmaps for genes within 20 kb of telomeres, were performed in-house using Python (pandas, seaborn).

### Whole-genome Sequencing (WGS)

WGS was performed by Novogene. Raw sequencing reads and processed data have been deposited in the NCBI Sequence Read Archive (SRA) under BioProject accession PRJNA1308857.

## Acknowledgment

This work was supported by NIH research grant GM152207 and American Cancer Society Research Scholar Award RSG-21-013-01-DMC to H.N. We thank the IU Bloomington Flow Cytometry Core Facility (RRID:SCR_024398) for their assistance with cell analysis. The CytoFLEX LX flow cytometer used in this study was funded in part by the IU Office of Research through the Research Equipment Fund. We thank Drs. Hannah Klein (New York University), Judith Campbell (Caltech), Patrick Sung (UT Health San Antonio) and Zhi Qi (Peking University) for providing yeast strains and plasmids, Kate Dinnon and Olivia Kavanaugh for technical support.

---

**Supplementary Figure 1.**
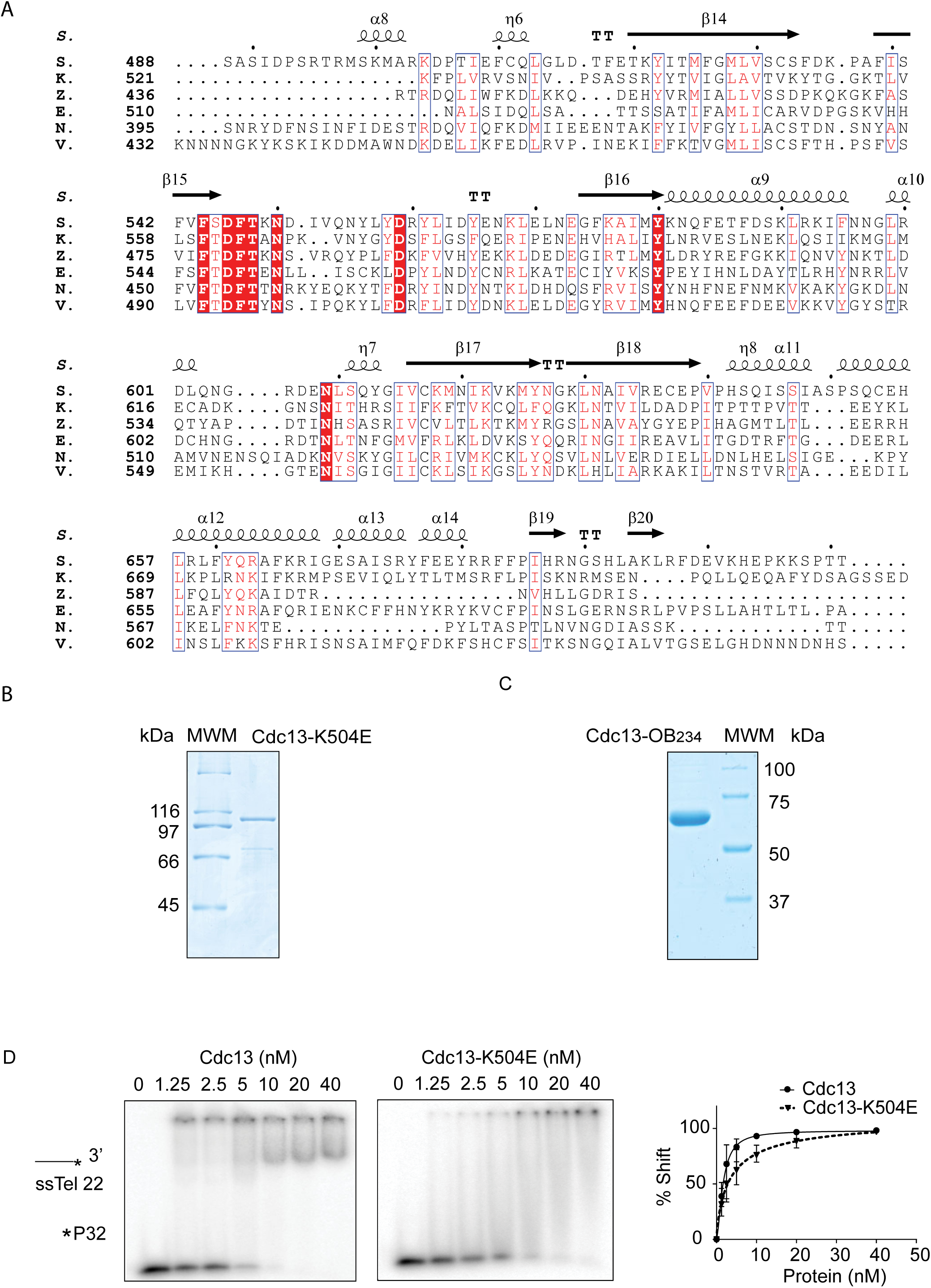
Expression, purification, and validation of Cdc13 protein. (A) Domain schematics of *S. cerevisiae* Cdc13. (B) SDS-PAGE analysis of SUMO-tagged Cdc13 purified from *E. coli*. The SUMO tag was cleaved prior to downstream experiments. (C) Size-exclusion chromatography (SEC) analysis of purified Cdc13 after SUMO tag removal. Elution was monitored by absorbance at 280 nm. Molecular weight standards (indicated above the elution profile) were used for column calibration. Cdc13 eluted at a position corresponding to ∼280 kDa, consistent with a dimeric species.

**Supplementary Figure 2.**
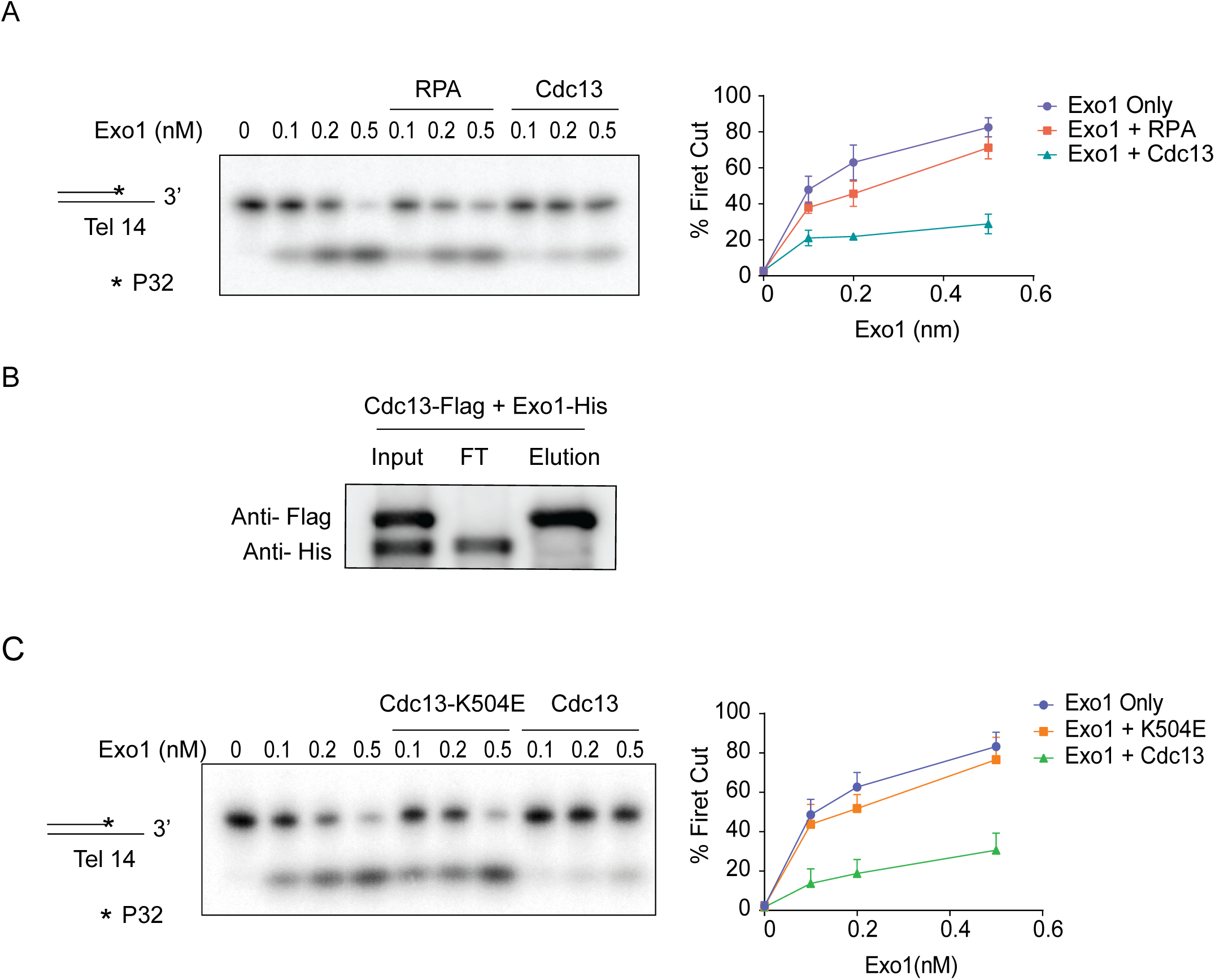
Biochemical characterization of Cdc13-K504E mutant. (A) Amino acid sequences of *K. lactis* and *S. cerevisiae* Cdc13 were aligned. Residues that are identical between the two species are highlighted in red, and residues with similar physicochemical properties are highlighted in pink. Alignment was performed using [Clustal Omega]. (B) SDS-PAGE analysis of SUMO-tagged Cdc13-K504E purified from *E. coli*. The SUMO tag was cleaved prior to downstream experiments. (C) SDS-PAGE analysis of SUMO-tagged Cdc13OB234 purified from *E. coli*. The SUMO tag was cleaved prior to downstream experiments. (D) EMSA comparing Cdc13 and Cdc13-K504E binding to a 5′- ^32^P-labeled ssTel22. Quantification curves represent mean ± SD from three independent replicates.

**Supplementary Figure 3.**
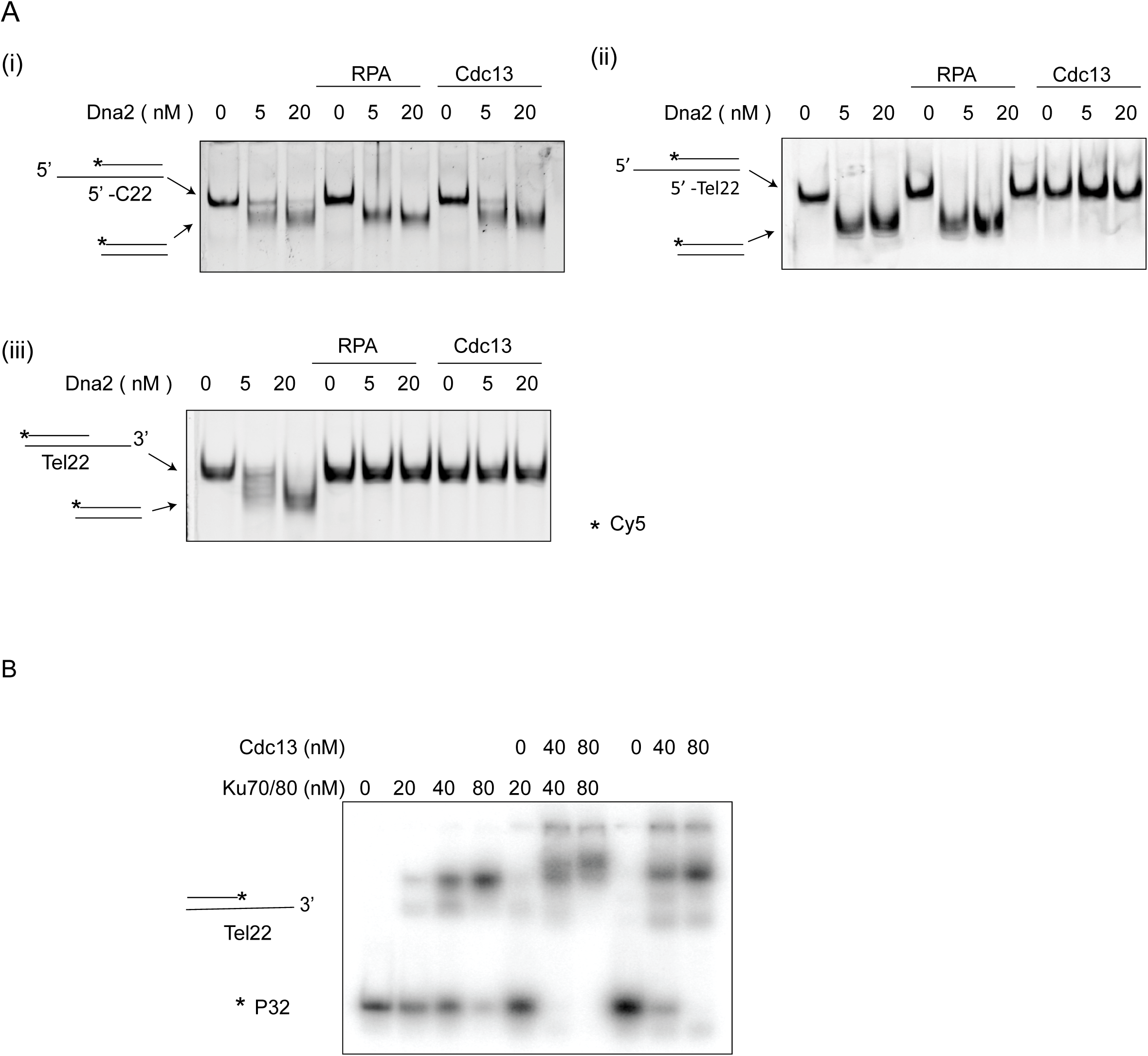
Control experiments for Exo1 resection analysis. (A) Cdc13, but not RPA significantly inhibits Exo1 resection on ^32^P-labeled Tel14. Quantification represents mean ± SD from three independent experiments. (B) Pull-down assay to assess interaction between Exo1 and Cdc13. (C) Cdc13, but not Cdc13-K504E significantly inhibits Exo1 resection on ^32^P-labeled Tel14. Quantification represents mean ± SD from three independent experiments.

**Supplementary Figure 4.**
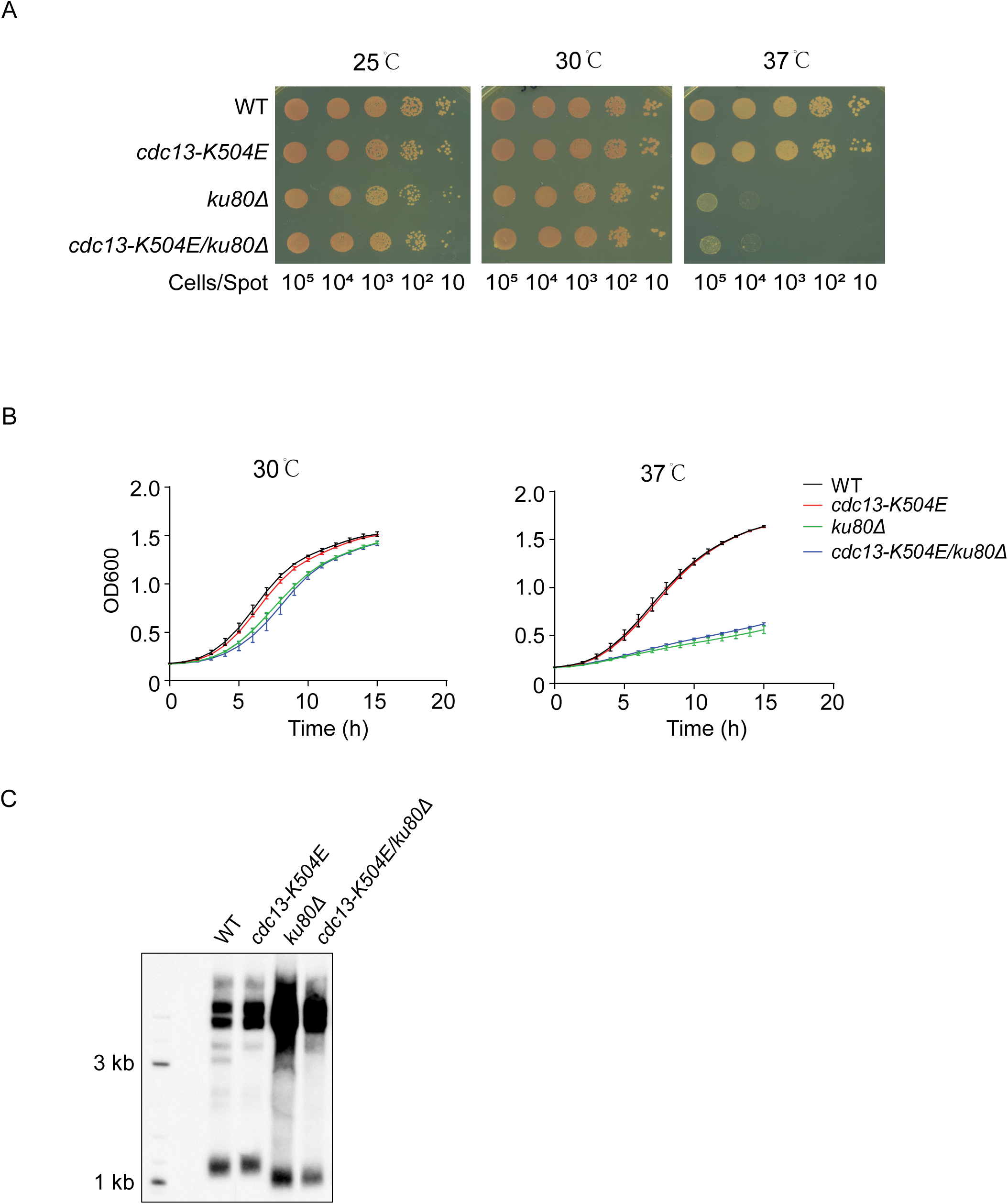
Control experiments for Sgs1-Dna2 resection analysis. (A) Dna2 nuclease assays on 5′-Tel22 (i), 5′-C22 (ii), and Tel22 (iii) substrates. All substrates were 3′ end–labeled with Cy5, as illustrated in the schematic. (B) Electrophoretic mobility shift assays (EMSA) of Ku and Cdc13 on the Tel22 substrate. Reactions contained Ku or Cdc13 alone, or both proteins added sequentially (Ku followed by Cdc13) to assess formation of the ternary Ku–Cdc13–DNA complex. Super-shifted complexes were detected.

**Supplementary Figure 5.**
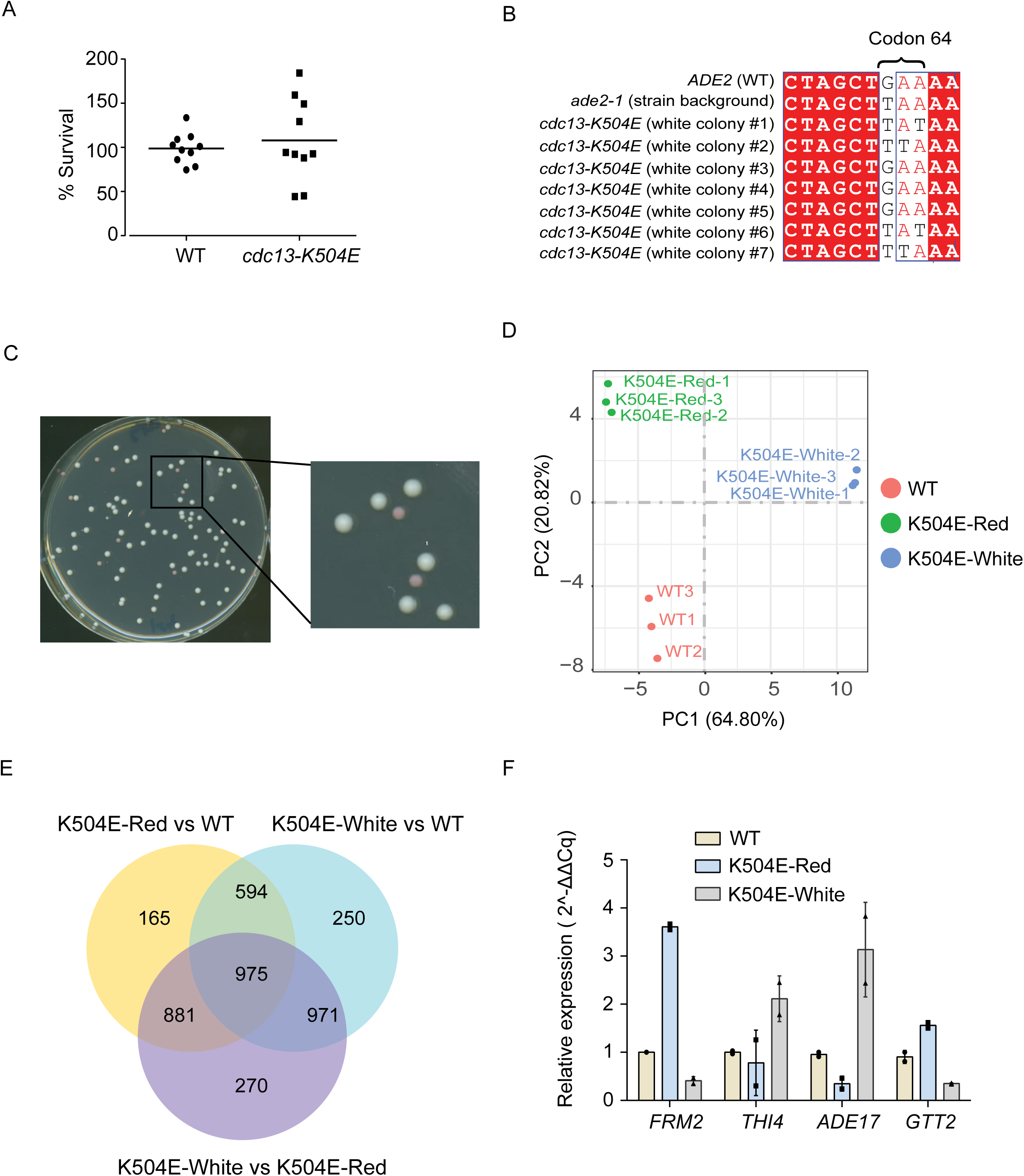
Additional in vivo phenotypic analyses of *cdc13*-K504E. (A) Temperature sensitivity assay of wild-type and *cdc13*-K504E strains with or without *ku80Δ* on YPD plates. Plates were incubated at 25°C, 30°C, or 37°C. (C) Growth curves of wild-type and *cdc13*-K504E strains with or without *ku80Δ* at 30°C and 37°C. (B) Southern blot analysis of telomere length in wild-type and *cdc13*-K504E strains with or without *ku80Δ* strains. (D) Variation assay of wild-type and *cdc13*-K504E strains. Each strain was plated on YPD medium after 15 days of growth. For each group, ten independent cultures were analyzed, and approximately 150 cells were plated per plate. (E) Sequence alignment of *ADE2* alleles from wild-type and *cdc13*-K504E revertant colonies. The *ADE2* locus was PCR-amplified and sequenced from one wild-type strain carrying the *ade2-1* mutation and seven white *cdc13*-K504E revertant colonies. The *ADE2* (WT) sequence was used as reference for alignment. (F) Inset of Figure 6B. Magnified view of the boxed region highlighting the size difference between white and red *cdc13*-K504E colonies.

**Supplementary Figure 6.**
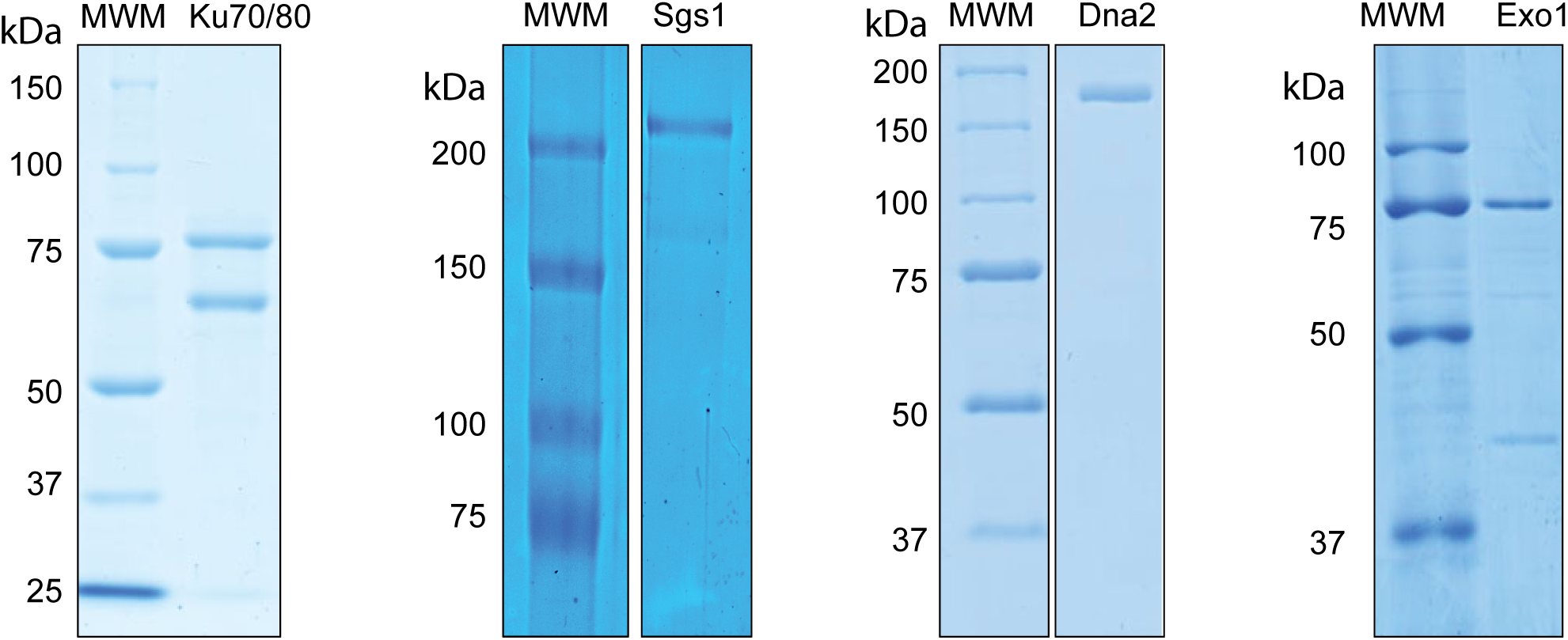
Analyses of *ADE2* revertants. (A) Growth curves of red and white colonies from wild-type and *cdc13*-K504E strains in liquid YPD. (B) Flow cytometry analysis of asynchronous cultures of WT, K504E-Red, and K504E-White strains. PI-Area (B610-A) vs Count histograms are shown. (C) Principal component analysis (PCA) of RNA-seq profiles from WT, K504E-Red, and K504E- White strains. (D) Venn diagram showing the overlap of differentially expressed genes among indicated comparisons. (E) RT-qPCR analysis of selected genes in WT, K504E-red, and K504E-white colonies. Relative mRNA levels were measured using Actin as the reference gene and normalized to WT-red. Data are presented as mean ± SD from two technical replicates.

**Figure S7.**
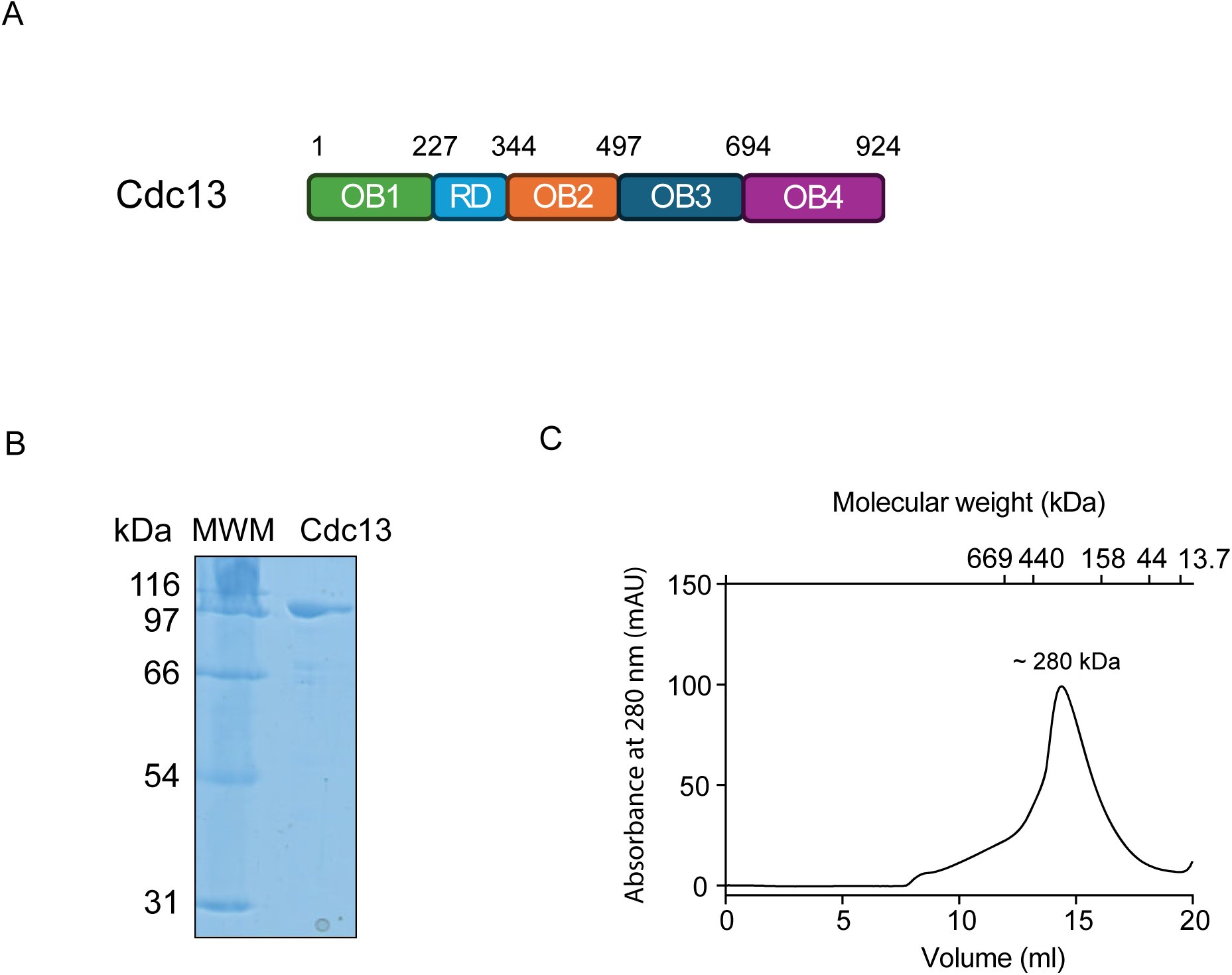
SDS-PAGE analysis of additional proteins used in this study. Coomassie-stained gels showing the indicated proteins. Molecular weight markers are included.

**Supplementary Figure S8.**
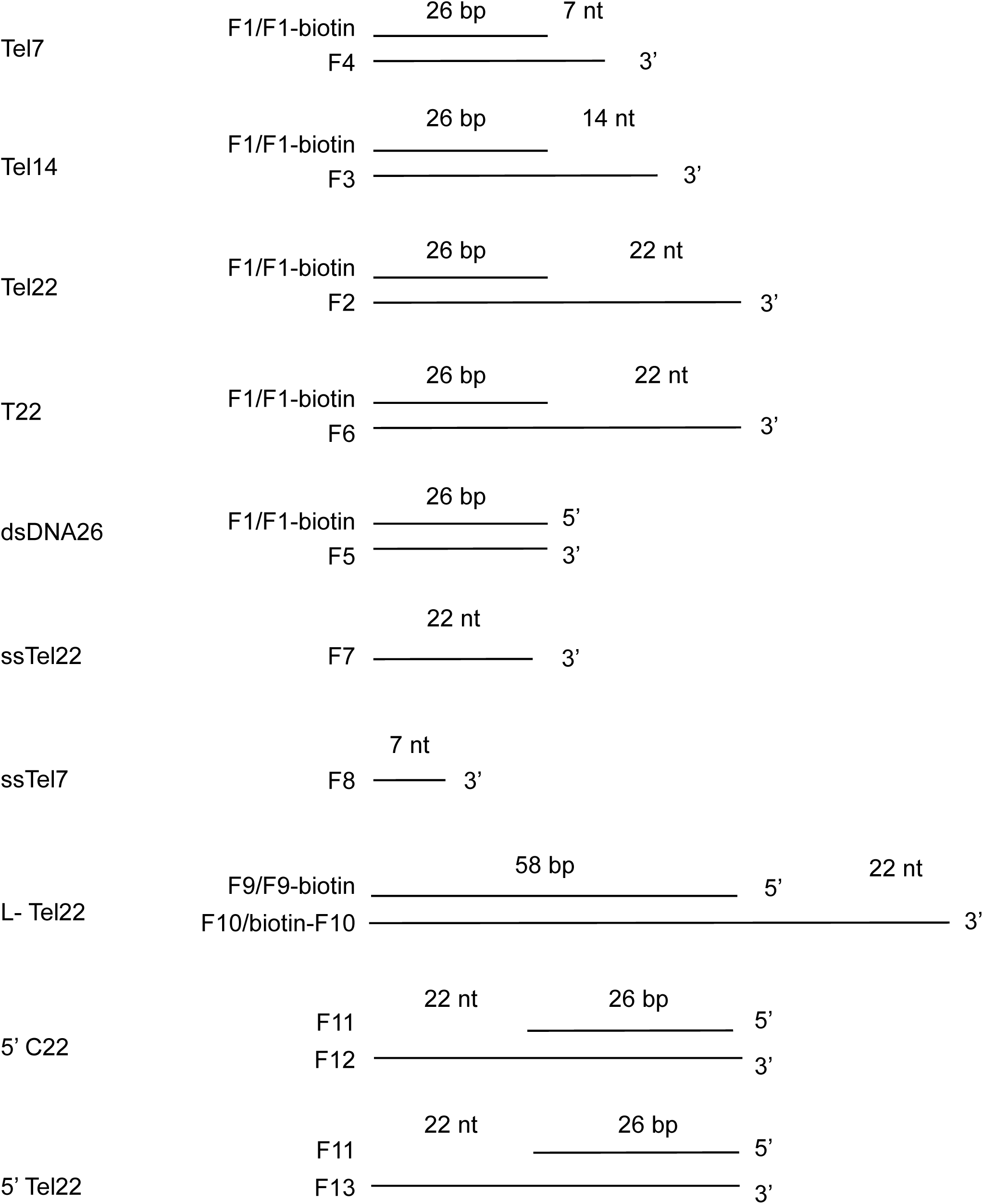
Summary of DNA substrates used in this study. Diagram summarizing the sequences, lengths, and labeling of DNA substrates employed in the experiments.

**Table S1.**
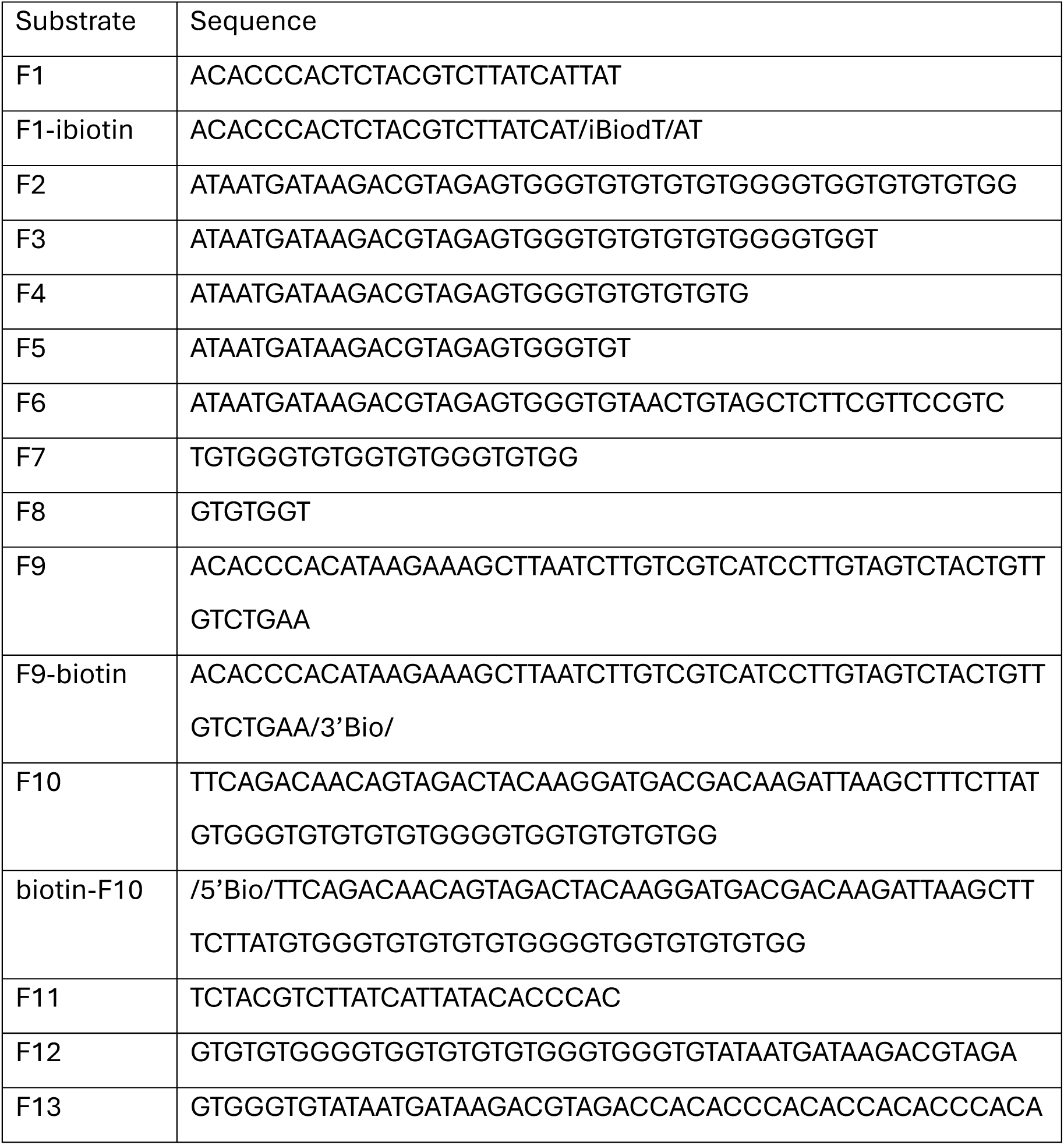
DNA substrates used in this study. List of DNA substrates, including sequences, lengths, and labeling information for all oligonucleotides used in experiments.

**Table S2.**
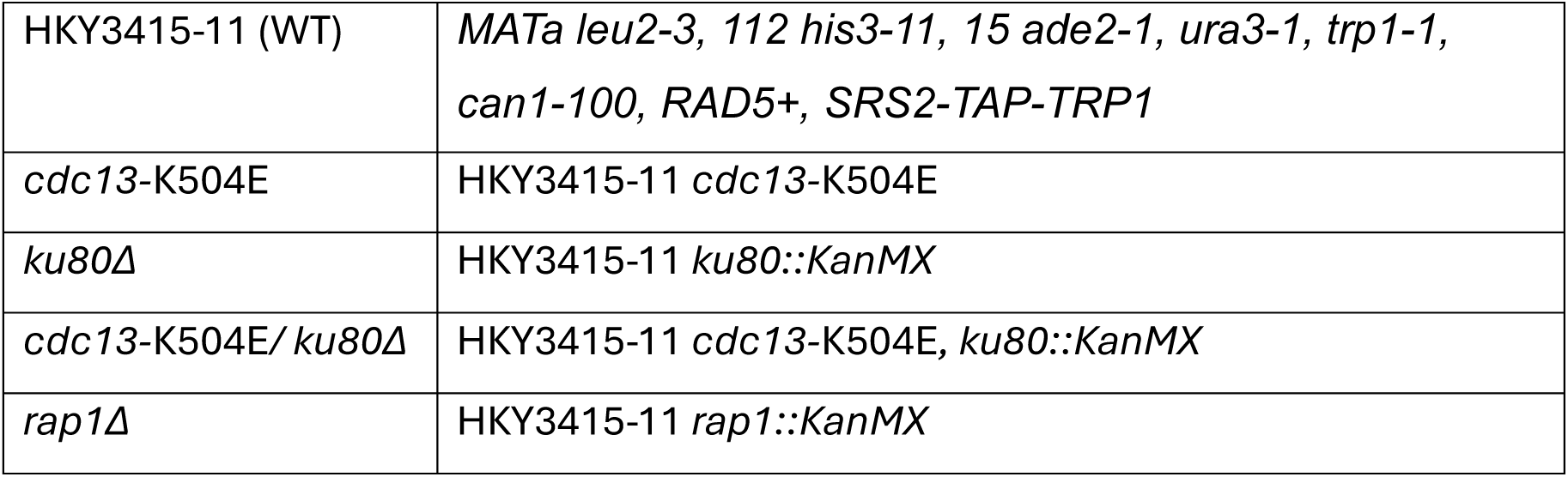
Yeast strains used in this study. Genotypes of all yeast strains employed in this study, including relevant mutations and background information.

## Notes

https://www.ncbi.nlm.nih.gov/bioproject/PRJNA1308857

### Competing Interest Statement

The authors have declared no competing interest.

### Summary of Updates

This version includes updated text and figures to reflect the latest analyses and clarifications made prior to journal submission. Minor edits were made throughout the manuscript for accuracy and clarity. Author affiliations were also updated.

